# 3MDR, a microcomputer-controlled visual stimulation device for psychotherapy-like treatments of mice

**DOI:** 10.1101/2022.11.10.516035

**Authors:** Isa Jauch, Jan Kamm, Luca Benn, Lukas Rettig, Hans-Christoph Friederich, Jonas Tesarz, Thomas Kuner, Sebastian Wieland

## Abstract

Post-traumatic stress disorder and other mental disorders can be treated by an established psychotherapy called Eye Movement Desensitization and Reprocessing (EMDR). In EMDR, patients are confronted with traumatic memories while they are stimulated with alternating bilateral stimuli (ABS). How ABS affect the brain and whether ABS could be adapted to different patients or mental disorders is unknown. Interestingly, ABS reduced conditioned fear in mice. Yet, an approach to systematically test complex visual stimuli and compare respective differences in emotional processing based on (semi-)automated behavioral analysis is lacking. We developed 3MDR (Model for MultiModal visual stimulation to Desensitize Rodents) - a novel, open-source, low-cost, customizable device that can be integrated in and TTL-controlled by commercial rodent behavioral setups. 3MDR allows to design and precisely steer multimodal visual stimuli in the head direction of freely-moving mice. Optimized videography allows to semi-automatically analyze rodent behavior during visual stimulation. Detailed building, integration, and treatment instructions along with open-source software provide easy access for inexperienced users. Using 3MDR, we confirmed that EMDR-like ABS persistently improve fear extinction in mice and showed for the first time that ABS-mediated anxiolytic effects strongly depend on physical stimulus properties such as ABS brightness. 3MDR not only enables researchers to interfere with mouse behavior in an EMDR-like setting, but demonstrates that visual stimuli can be used as a noninvasive brain stimulation to differentially alter emotional processing in mice.

**SIGNIFICANCE STATEMENT:** Alternating bilateral stimuli (ABS) reduce fear in post-traumatic stress disorder patients and in mice. The mechanism of how classic ABS – typically used in Eye Movement Desensitization and Reprocessing (EMDR) - reduce fear is enigmatic. We provide detailed resources to build a cost-effective, computer-controlled device called 3MDR to perform and semi-automatically analyze EMDR-like treatments in freely-moving mice and to test behavioral effects of multiple ABS variants. Using the 3MDR device, this study confirmed that classic ABS strongly and persistently improve the extinction of conditioned fear in mice – an effect that depended on the brightness of ABS. This novel method may ultimately contribute to a deeper translational and neurobiological understanding of how visual stimuli affect emotional processing in mice.

## INTRODUCTION

Alternating bilateral stimuli (ABS) are the key component of an established psychotherapy against post-traumatic stress disorder (PTSD) called Eye Movement Desensitization and Reprocessing (EMDR) (Shapiro, 1989; World Health Organization and Van Ommeren, 2013). In EMDR, a patient is recalling a traumatic memory while being exposed to classic ABS. Classic ABS are defined in that the patient shifts the gaze in order to pursue a visual stimulus such as a light beam or the therapist’s hand that horizontally oscillates at a stable frequency of ∼1 Hz. Combining exposure with classic ABS improves extinction of traumatic memories and reduces hyperarousal, vividness of memories and intrusions (Calancie et al., 2018; Rousseau et al., 2019). EMDR has been adapted to many mental disorders, e.g. chronic pain, depressive or anxiety disorders, through disorder-specific exposure protocols (Fereidouni et al., 2019; Feske and Goldstein, 1997; Tesarz et al., 2014, 2013) but not through significant adaptations of the classic ABS paradigm. Despite its clinical evidence, it remains unresolved how ABS alter emotional processing in patients (Calancie et al., 2018). A better mechanistic understanding would provide insights into whether ABS variants could further improve EMDR treatments of specific disorders or individuals.

Human research suggests that EMDR differs from conventional exposure therapies as ABS directly affect the processing of traumatic memories (Lee and Cuijpers, 2013). A recent animal study revealed a deeper neurobiological understanding of ABS-mediated effects (Baek et al., 2019): This animal model of EMDR required a coincident presentation of both ABS and the fear-associated cue for therapeutic benefits and revealed an ABS-mediated activation of a novel circuit in the superior colliculus (SC) - known for multisensory integration and saccade control-that suppressed cue-related fear engrams in amygdala.

An animal model of EMDR allows to investigate interactions between ABS and the brain not only on the level of brain circuits but also on a procedural level of respective ABS properties. With more options in terms of standardization and feasibility compared to human studies, a suitable EMDR-like animal model could elucidate which ABS properties causally mediate anxiolytic effects in mice and which ABS variant optimally reduces fear. To investigate the influence of physical ABS properties on mice, researchers need a system capable of creating a plethora of stimulus qualities and enable standardized and EMDR-like experiments in mice. Unfortunately, there are no commercial systems available to perform such rodent experiments.

We successfully built a customizable, low-cost device allowing the experimenter to design and center various visual stimuli in the head direction of freely-moving mice. We share detailed instructions on hardware and software to ease the setup process for inexperienced researchers. Furthermore, we validated our system and confirmed that classic ABS persistently improve fear extinction in mice and showed for the first time that these anxiolytic effects depend on specific physical ABS properties, such as ABS brightness. This new method has the potential to substantially improve the study of EMDR and of noninvasive visual brain stimulation in mice.

## MATERIALS AND METHODS

### System overview and integration

3MDR, a Model for MultiModal stimulation to Desensitize Rodents, consists of an Arduino-based microcontroller, a treatment cylinder, a remote control and infrared videography based on an infrared floor illumination (Fig. 1A;B). The main electronic component of the 3MDR apparatus (Table 2) is an Arduino Uno microcontroller board (https://www.arduino.cc, Fig. 2A, Fig. 1-1A), which is programmed with a custom script that encodes ABS stimulus characteristics. We provide four baseline scripts to generate either horizontally, vertically or two different patterns of oppositely oscillating visual stimuli to investigate anxiolytic effects of different types of stimulation (Video 1). All four scripts work in an analogous fashion and are commented for users without coding expertise (Fig. 2-1). The following parameters can easily be defined in our scripts: stimulus width (number of LEDs used), alternation frequency (Hz), stimulus color and brightness, stimulus duration after trigger and row pattern. The digital output signals of the Arduino Uno communicate with up to nine LED strips (Fig.1D, 2C, APA104, Sparkfun, USA) via general purpose input/output (GPIO) pins (Fig. 1-1B). Each LED strip contains 36 LEDs under a waterproof silicone shield (Fig. 1-1D, E). These LED strips are attached to the walls of a black vinyl or acrylic cylinder, in which the visual stimuli can be presented to the mice. The number and alignment of individual LED strips can be flexibly chosen for every setup to perform experiments with different visual stimulus patterns. We recommend to cover the inside of the cylinder with a non-reflective, resistant coating, e.g. ultramat 2-component paint (Crop 2K Spraymax, RAL 9005, ultramat, Netherlands) or alternatives with transparent ultramat paint (Aerodecor transparent matt, Modulor, Germany) to carefully prevent light scattering during ABS application (Fig. 1C).

**Figure 1:**
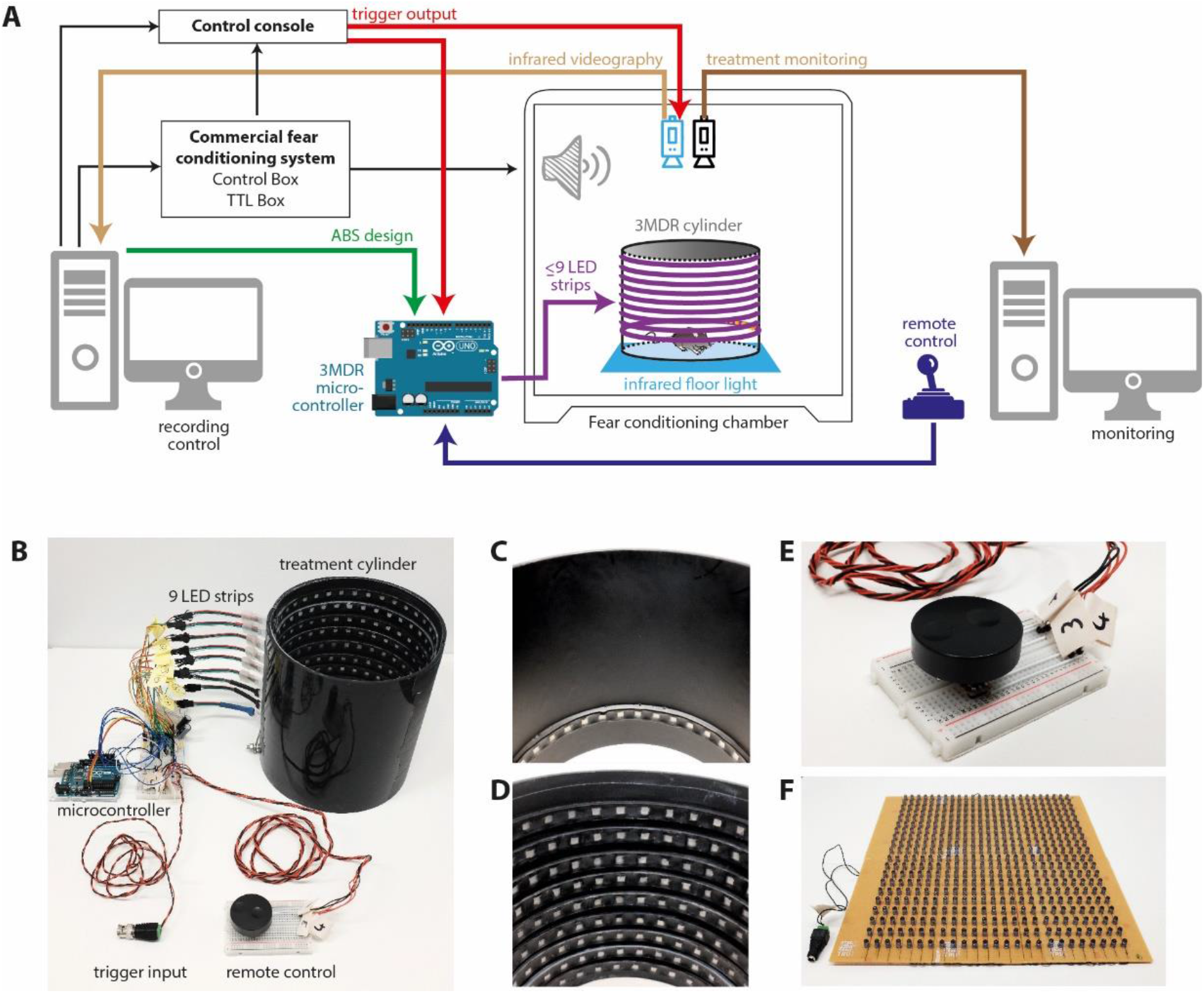
Overview and integration of 3MDR system in a fear conditioning setup. A) Schematic overview how to integrate 3MDR in a commercially available fear conditioning system. B-E) Photographs of the 3MDR system. B) Overview showing all electronic components including the remote control, trigger input and treatment cylinder. C-D) Two example configurations of the treatment cylinder with a single (C) and 9 rows (D) of LED strips. E) Closeup image of the remote control. F) Low construction of infrared floor illumination panel made of a 25×20 LED array.

**Figure 2:**
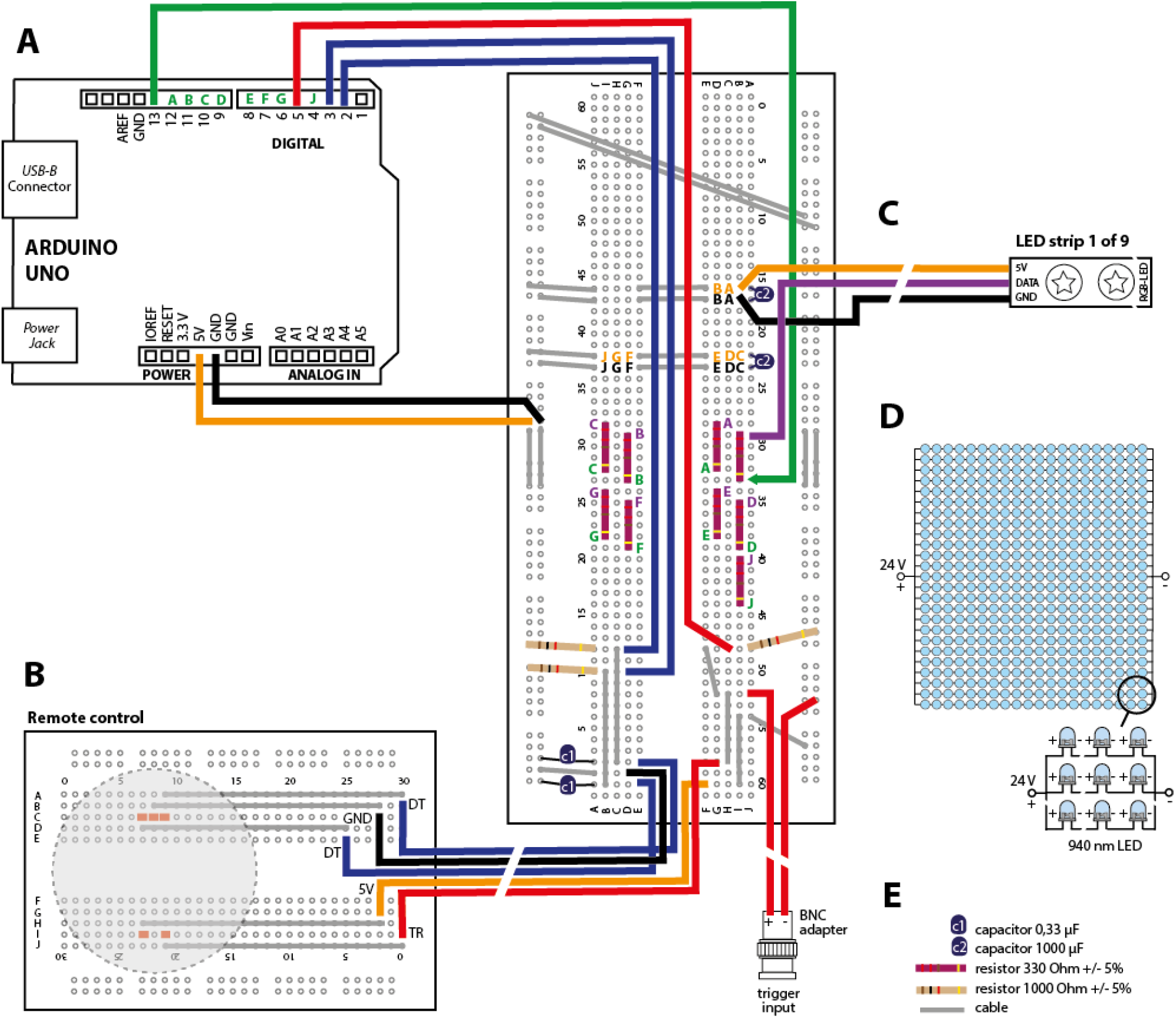
3MDR hardware components. A schematic showing the hardware part of 3MDR based on an Arduino Uno microcontroller (A). GPIO pins 2 and 3 receive rotation data (dark blue) of remote control (B). Arduino’s GPIO pins 4, 6-13 convey data of ABS design and movement (green) to control LED strips (C). GPIO pin 5 receives trigger information (red) from external trigger input or manual trigger input from remote control. We displayed the connection principle of one of the nine LED strips (LED strip H) whereas the connections of the other eight LED strips are color-coded for clarity. Each LED-strip (A-J) needs 4 cable connections (green pins: connection between board and Arduino Uno. violet pins: connection between board and LED strip data input; orange pins: 5 V supply; black pin: ground circuit). D) A schematic of our infrared floor light showing a parallel circuit of 25 rows of 20 LEDs (940 nm wavelength). E) Visual figure legend

The stimulus presentation is initiated by an electrical input signal which can be triggered via the GPIO pin to the Arduino Uno either manually or electronically Manual input trigger signals can be delivered by a simple push-button operation at the remote control (Fig. 1E, 2B; Fig. 1-1C). Electronic input trigger signals, e.g. generated by TTL output boards of commercial behavioral setups, can be connected to the Arduino Uno via BNC connection. This enables users to easily integrate the apparatus into existing rodent behavioral setups. We used our 3MDR system with a regular auditory fear conditioning system (Maze Engineers, USA) and a TTL trigger unit (Maze Engineers, USA). We isolated the fear conditioning box with acoustic foam (Flat Tec 2 cm, AixFoam, Germany) to optimize sound presentation. A custom-made infrared floor illumination panel based on 940 nm LEDs (Fig. 1F, 2D; Table 3) was placed under a semitransparent acrylic floor to increase the visual contrast of mouse behavior in video recordings and to facilitate behavioral analyses. We integrated two top-view cameras in the fear conditioning box above the conditioning chamber: An infrared camera (DMK 37BUX265 and lens TPL 0620 6MP, Imaging Source, Germany; 850 nm longpass filter, FEL0850, Thorlabs, Germany) recorded animal behavior. Freezing behavior was quantified in infrared videos based on detection of frame-to-frame pixel changes (Pennington et al., 2019). A second treatment monitoring camera (Logitech HD webcam C310) enabled the experimenter to visualize and manually center the position of ABS in head direction of mice during fear extinction (Video 2). Dynamic 360° shifting is achieved by a rotary encoder which communicates with the Arduino Uno via two GPIO pins (Fig. 2B). The fear conditioning chamber, camera recording and 3MDR apparatus were synchronized in a master-slave architecture: the TTL output unit of the fear conditioning box generated TTL signals under the control of original software (Maze Origin, Maze Engineers, USA). TTL signals were fed in a control console (Fiber photometry console, Doric Neuroscience, Canada) and visualized in its original software (Doric Neuroscience Studio, Canada). Consecutively, the control console triggered the infrared video recordings and synchronized CS and ABS presentations. Before the start of ABS presentations, the experimenter chose, adjusted and uploaded the suitable Arduino script on the microcontroller. TTL-input to the Arduino Uno (https://www.arduino.cc) triggered the preset ABS presentation within the experiment.

### Animals

In the study, 11-17 weeks old male C57BL6 wild-type mice were used, which were housed in small groups (maximum 4 in one cage of an IVC system) with food and water *ad libitum* under a 12 h :12 h light-dark cycle (light off at 7:00 am). All experiments were carried out during the dark phase of the cycle. Two days before the start of the experiment, male mice were single-housed to reduce between-subject influences of ABS treatment. All animal procedures were performed in accordance with the (Author University) animal care committee’s regulations. Efforts were made to minimize animal suffering and to reduce the number of animals used according to the 3R’s principles. Experimenters were blinded to the identity of the mice during the analysis of behavioral tests.

### Behavioral experiments

We used our 3MDR system in combination with a fear conditioning setup during fear extinction experiments in mice (Curzon et al., 2009). Fear behavior was quantified as freezing times.

#### Habituation

Mice were transferred to a holding room neighboring the experimental room and rested for 1 hour before and after experiments. For three consecutive days, mice were handled by the experimenter for a few minutes and habituated in context A for 15 min per day. Before each experiments, mice were shortly habituated for 120 s in the respective contexts.

*Auditory fear conditioning* (FC, day 1) was performed in context A by pairing a conditioned stimulus (CS, 3 kHz tone, 30 s continuous, 75 dB) either (classic paradigm) four times with a soft unconditioned stimulus (US1, electric foot shock, 0.4 mA, 1 s) or (IFE paradigm) five times with a stronger electric foot shock (US2, 0.7 mA, 1 s). CS and US co-terminated and CS-US combinations were presented at pseudo-randomized inter-trial intervals (ITIs) between 90 to 150 s. We excluded animals that did not sufficiently learn the CS-US association showing maximum freezing times below 65% CS time (5 out of 30 animals).

#### Fear extinction (FE)

On day 2, we presented CS 20 times at pseudo-randomized ITIs of 60-90 s and measured CS-mediated freezing behavior in context B. In control groups, CS was presented without ABS. In treatment groups, ABS or XBS were paired with CS from the second CS presentation onwards.

#### Fear recall (FR)

On day 10, we presented 3 CS at pseudo-randomized ITI of 90-150 s in context B and measured CS-mediated freezing behavior.

#### Fear renewal (FN)

At least two hours after recall tests on day 10, we presented 3 CS at pseudo-randomized ITI of 90-150 s in a novel context C.

#### Experimental environments during fear conditioning

To reduce the degree of contextual fear associations, behavioral experiments were performed in different contexts: Context A consists of a square experimental chamber (16 cm * 16.5 cm * 25 cm) with grey monochrome walls, a shock grid with metal rods (Maze Engineers, USA) and white light illumination (30 Lux). Context B is the round 3MDR cylinder (diameter 20 cm, height 20 cm) on a smooth acrylic glass base plate in no-visible-light conditions and scented with 1,4-cineol (Sigma Aldrich, USA). Context C consists of a square chamber (16 cm * 16.5 cm * 25 cm) with black and white striped walls (Maze engineers, USA), a textured, custom base floor plate, white light illumination (7 Lux) and a rose blossom scent (scented creme, Rituals). All contexts were continuously illuminated by our infrared floor light panel.

### ABS treatment during fear extinction

ABS features were preset in our custom-made Arduino script and uploaded to the microcontroller before the start of the experiment (Fig. 2-1). We used the following settings for classic ABS (Fig. 4): In the Arduino script “Full_cylinder_horizontal_with_rotary_encoder”, we adjusted the oscillation length = 9 (number of LEDs used), alternation frequency = 1 (Hz), stimulus color and brightness Red = 30, Green = 30, Blue = 30, stimulus duration = 30 s and activated LED strip 2. Extrabright bilateral stimuli (Figure 5) consisted of increased channel values for Red = 255, Green = 255 and Blue = 255. The duration of both CS and ABS was 30 s and both ABS and CS started simultaneously. TTL signals automatically started ABS presentation that consisted of an oscillating light beam in which nine LEDs were switched on serially from left to right and vice versa with an alternation frequency of 1 Hz.

### Freezing analysis

Infrared videography was recorded at a framerate of 25 frames per second. Infrared video material enabled us to semi-automatically score freezing behavior based on frame-to-frame pixel change with the open-source algorithm ezTrack (Pennington et al., 2019). The motion threshold and animal’s freezing threshold were defined as described in Pennington et al‥ We scored freezing if pixel changes (a correlate of animal movements) were below the freezing threshold for at least 25 consecutive frames (=1 s). We analyzed the freezing time during CS presentation and scored freezing as a percentage of CS-time (Freezing (%) = freezing time (s)/30 s). We initially validated whether infrared and visible light videography enables comparable freezing detection during ABS presentation (Fig. 2-2). We simultaneously recorded animal behavior with our monitoring and infrared camera (Video 3) during two CS-ABS presentations and performed semi-automated freezing analysis on both types of videographic recordings (Fig. 2-2A, B). Infrared recordings facilitated the recording of freezing behavior throughout ABS stimulation, whereas visible light videography diminishes pixel change analysis during ABS treatments due to massive light artifacts (paired t-test, n = 12 CS (6 mice), p = 0.0^a^, Fig. 2-2D, Fig. 2C).

### Statistics

Statistical analyses were performed using GraphPad Prism 6 and www.estimationstats.com. Statistical significance was defined as a value of p < 0.05. Data are presented as mean +/-SEM if not mentioned differently. Population size (n) is presented as number of mice if not mentioned differently. We tested our datasets for deviations from normality with the D’ Agostino-Pearson Test and Shapiro-Wilk test.

Comparisons between more than two groups were analyzed by repeated-measures two-way ANOVA followed by Sidak’s test for multiple comparisons or One-way ANOVAs followed by Tukey’s test for multiple comparison (Table 1). Comparisons between two groups (Fig. 4-6) were computed in www.estimationstats.com and displayed in Gardner-Altman estimation plots. For two paired groups, we computed the paired Hedges’ g and displayed a slopegraph. For two independent groups, we computed the Cliff’s delta and displayed a scatter dot plot. The respective mean differences were plotted on floating axes on the right as a bootstrap sampling distribution. The mean difference is depicted as a dot; the 95% confidence interval is indicated by the ends of the vertical error bar. 5000 bootstrap samples were taken; the confidence interval is bias-corrected and accelerated. We computed P values of two-sided permutation t-tests. The P value(s) reported are the likelihood(s) of observing the effect size(s), if the null hypothesis of zero difference is true. For each permutation P value, 5000 reshuffles of the control and test labels were performed. To compare the strength of US-CS association between groups, we analyzed the maximal freezing response (within all FC sweeps of each animal) during FC test (referred to as FC max). To investigate the strength of fear extinction, we computed the average freezing responses during the last three FE sweeps per animal (referred to as FE end).

**Table 1:**
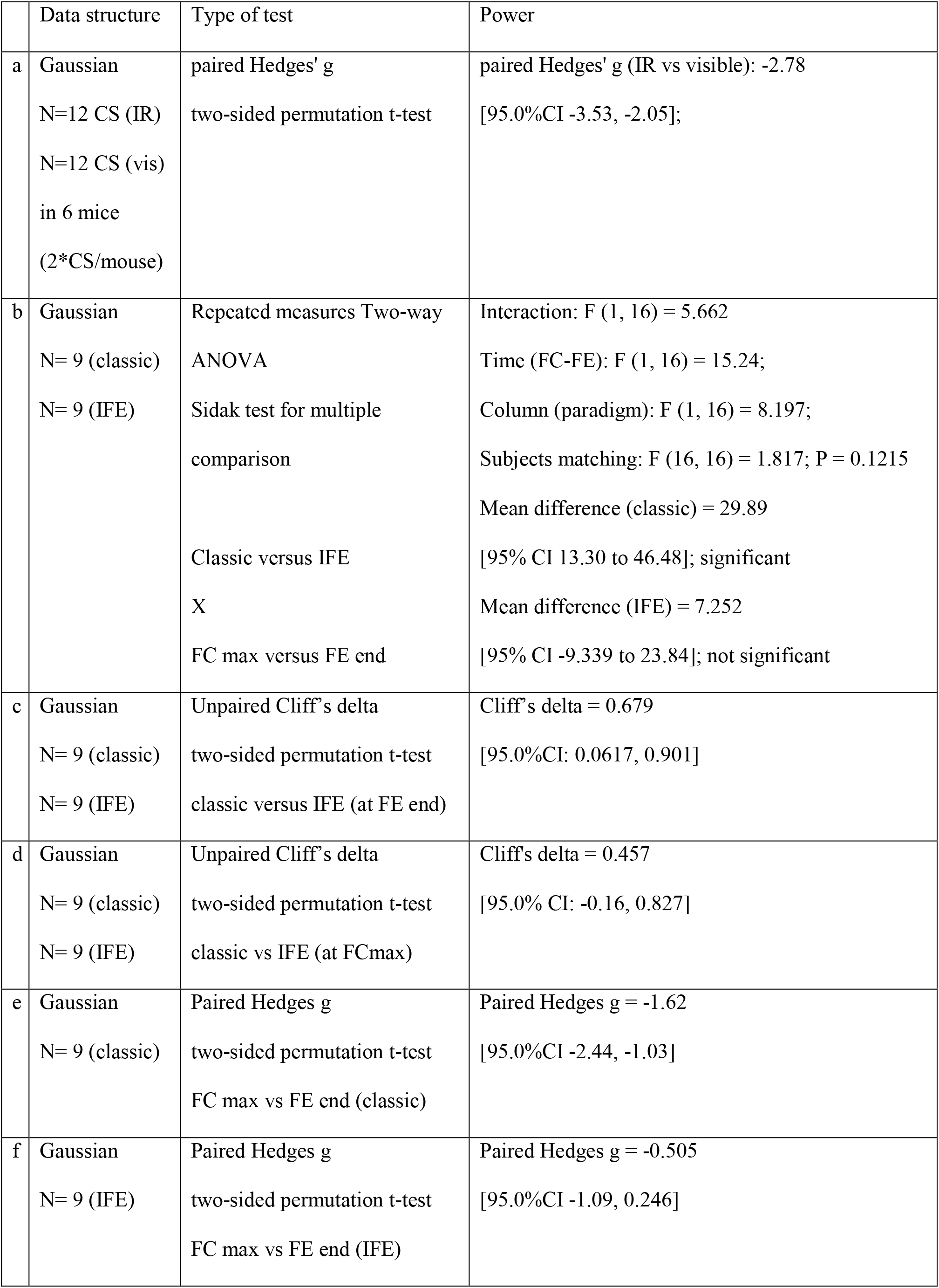
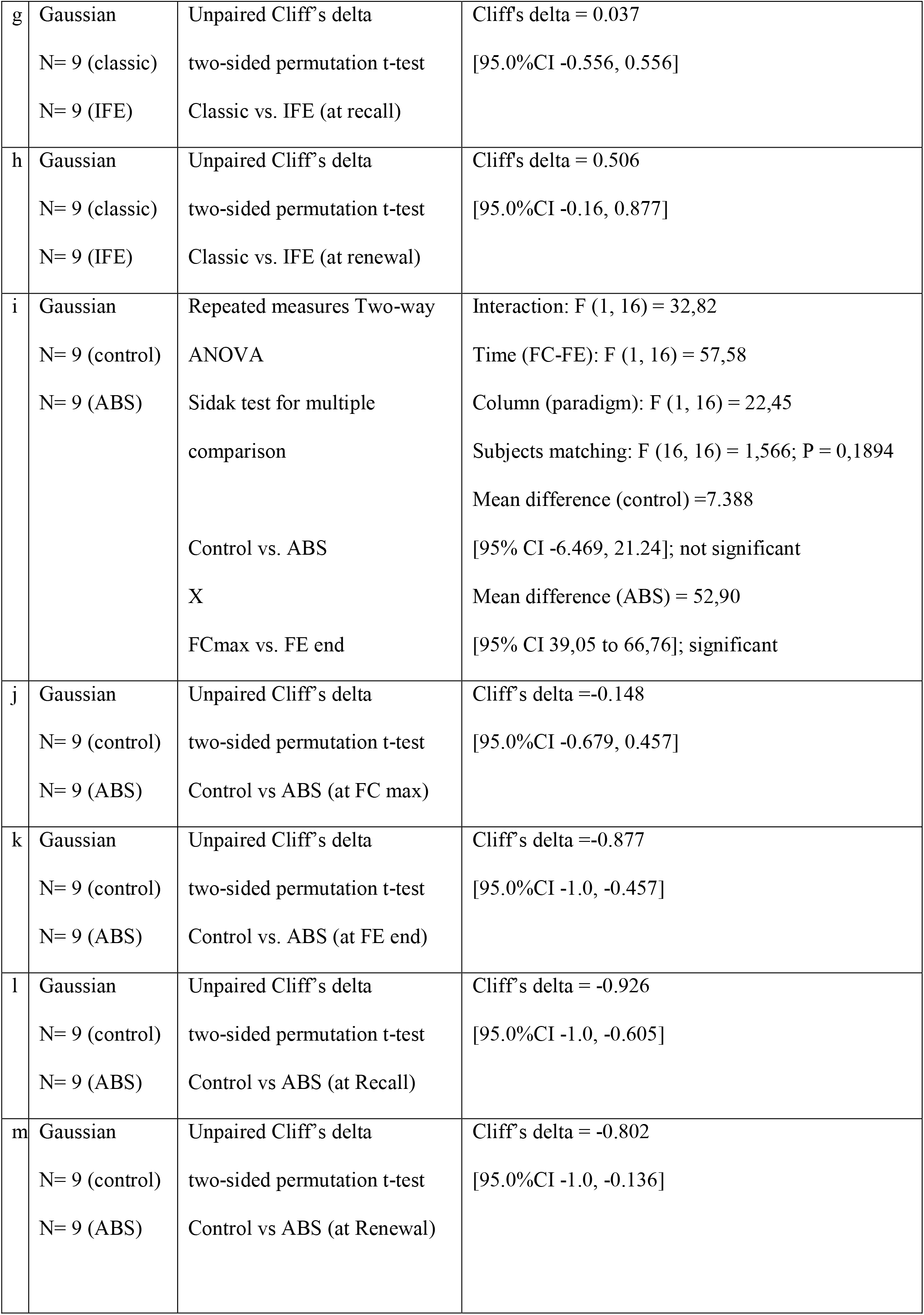
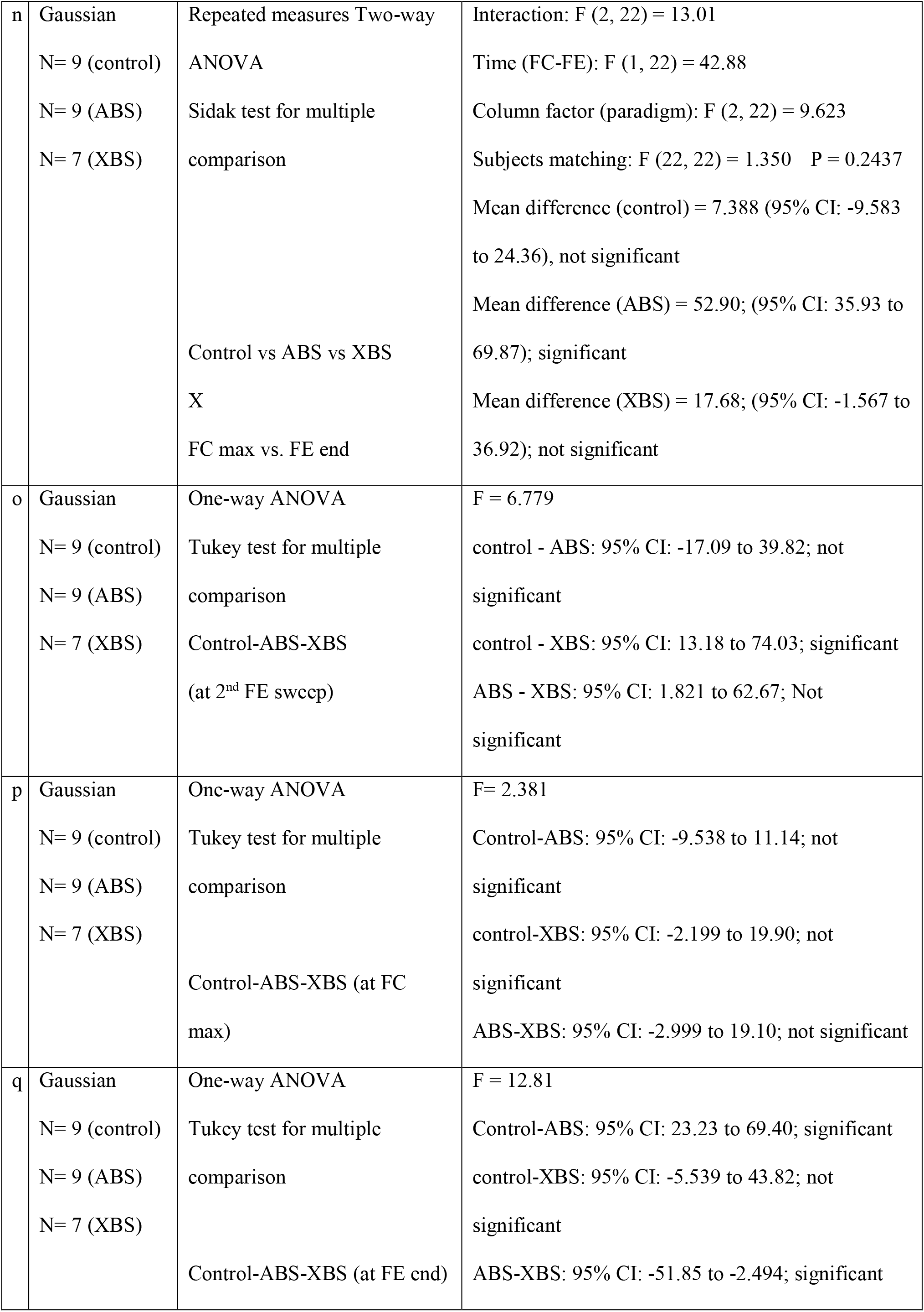
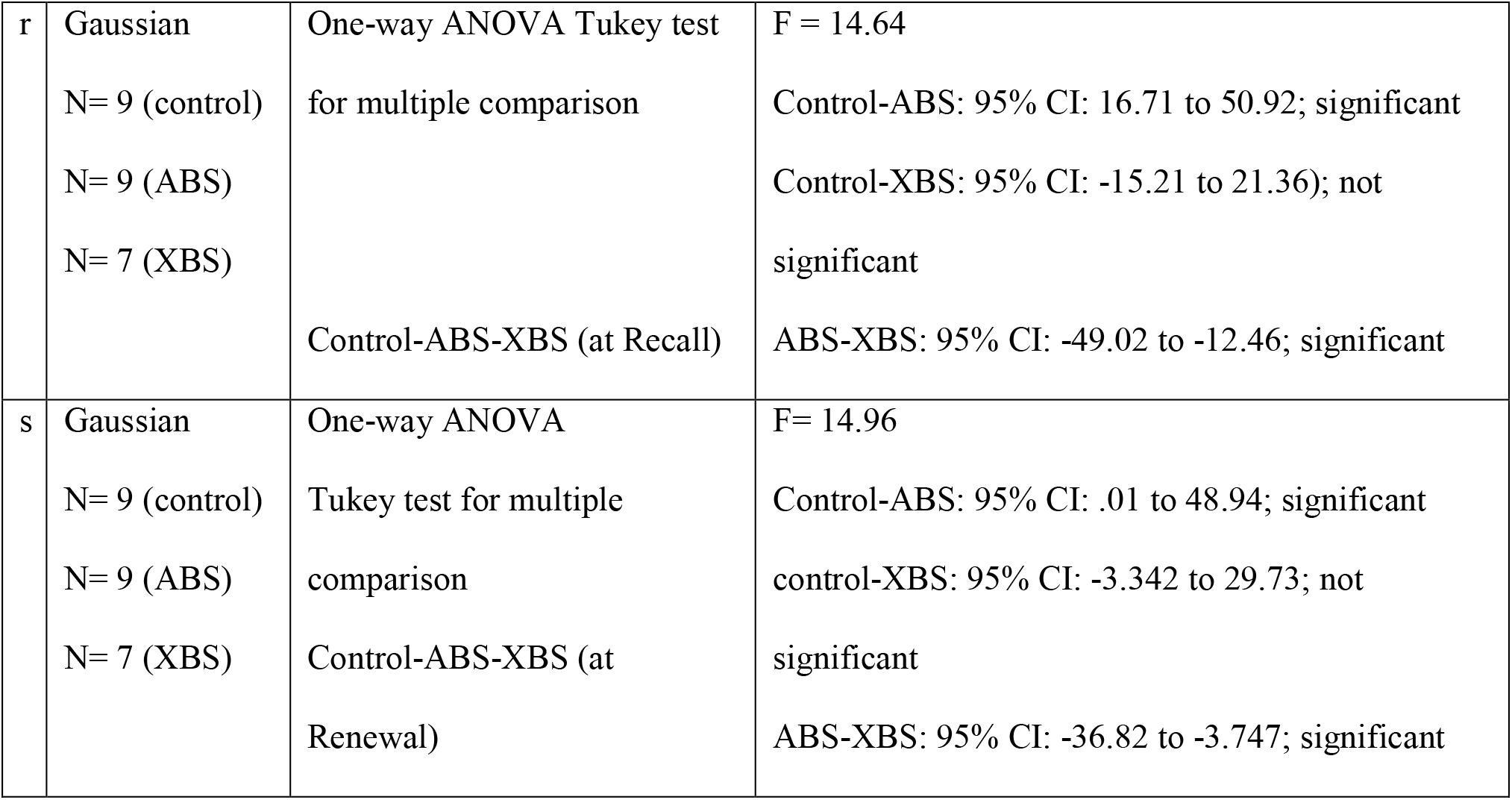
Statistical table.

|  | Data structure | Type of test | Power |
| --- | --- | --- | --- |
| a | Gaussian<br>N=12 CS (IR)<br>N=12 CS (vis)<br>in 6 mice<br>(2*CS/mouse) | paired Hedges' g<br>two-sided permutation t-test | paired Hedges' g (IR vs visible): -2.78<br>[95.0%CI -3.53, -2.05]; |
| b | Gaussian<br>N= 9 (classic)<br>N= 9 (IFE) | Repeated measures Two-way<br>ANOVA<br>Sidak test for multiple<br>comparison<br>Classic versus IFE<br>X<br>FC max versus FE end | Interaction: F (1, 16) = 5.662<br>Time (FC-FE): F (1, 16) = 15.24;<br>Column (paradigm): F (1, 16) = 8.197;<br>Subjects matching: F (16, 16) = 1.817; P = 0.1215<br>Mean difference (classic) = 29.89<br>[95% CI 13.30 to 46.48]; significant<br>Mean difference (IFE) = 7.252<br>[95% CI -9.339 to 23.84]; not significant |
| c | Gaussian<br>N= 9 (classic)<br>N= 9 (IFE) | Unpaired Cliff's delta<br>two-sided permutation t-test<br>classic versus IFE (at FE end) | Cliff's delta = 0.679<br>[95.0%CI: 0.0617, 0.901] |
| d | Gaussian<br>N= 9 (classic)<br>N= 9 (IFE) | Unpaired Cliff's delta<br>two-sided permutation t-test<br>classic vs IFE (at FCmax) | Cliff's delta = 0.457<br>[95.0% CI: -0.16, 0.827] |
| e | Gaussian<br>N= 9 (classic) | Paired Hedges g<br>two-sided permutation t-test<br>FC max vs FE end (classic) | Paired Hedges g = -1.62<br>[95.0%CI -2.44, -1.03] |
| f | Gaussian<br>N= 9 (IFE) | Paired Hedges g<br>two-sided permutation t-test<br>FC max vs FE end (IFE) | Paired Hedges g = -0.505<br>[95.0%CI -1.09, 0.246] |
| g | Gaussian<br>N= 9 (classic)<br>N= 9 (IFE) | Unpaired Cliff's delta<br>two-sided permutation t-test<br>Classic vs. IFE (at recall) | Cliff's delta = 0.037<br>[95.0%CI -0.556, 0.556] |
| h | Gaussian<br>N= 9 (classic)<br>N= 9 (IFE) | Unpaired Cliff's delta<br>two-sided permutation t-test<br>Classic vs. IFE (at renewal) | Cliff's delta = 0.506<br>[95.0%CI -0.16, 0.877] |
| i | Gaussian<br>N= 9 (control)<br>N= 9 (ABS) | Repeated measures Two-way<br>ANOVA<br>Sidak test for multiple<br>comparison<br>Control vs. ABS<br>X<br>FCmax vs. FE end | Interaction: F (1, 16) = 32,82<br>Time (FC-FE): F (1, 16) = 57,58<br>Column (paradigm): F (1, 16) = 22,45<br>Subjects matching: F (16, 16) = 1,566; P = 0,1894<br>Mean difference (control) = 7.388<br>[95% CI -6.469, 21.24]; not significant<br>Mean difference (ABS) = 52,90<br>[95% CI 39,05 to 66,76]; significant |
| j | Gaussian<br>N= 9 (control)<br>N= 9 (ABS) | Unpaired Cliff's delta<br>two-sided permutation t-test<br>Control vs ABS (at FC max) | Cliff's delta = -0.148<br>[95.0%CI -0.679, 0.457] |
| k | Gaussian<br>N= 9 (control)<br>N= 9 (ABS) | Unpaired Cliff's delta<br>two-sided permutation t-test<br>Control vs. ABS (at FE end) | Cliff's delta = -0.877<br>[95.0%CI -1.0, -0.457] |
| l | Gaussian<br>N= 9 (control)<br>N= 9 (ABS) | Unpaired Cliff's delta<br>two-sided permutation t-test<br>Control vs ABS (at Recall) | Cliff's delta = -0.926<br>[95.0%CI -1.0, -0.605] |
| m | Gaussian<br>N= 9 (control)<br>N= 9 (ABS) | Unpaired Cliff's delta<br>two-sided permutation t-test<br>Control vs ABS (at Renewal) | Cliff's delta = -0.802<br>[95.0%CI -1.0, -0.136] |
| n | Gaussian<br>N= 9 (control)<br>N= 9 (ABS)<br>N= 7 (XBS) | Repeated measures Two-way ANOVA<br>Sidak test for multiple comparison<br>Control vs ABS vs XBS<br>X<br>FC max vs. FE end | Interaction: $F(2, 22) = 13.01$<br>Time (FC-FE): $F(1, 22) = 42.88$<br>Column factor (paradigm): $F(2, 22) = 9.623$<br>Subjects matching: $F(22, 22) = 1.350$ $P = 0.2437$<br>Mean difference (control) = 7.388 (95% CI: -9.583 to 24.36), not significant<br>Mean difference (ABS) = 52.90; (95% CI: 35.93 to 69.87); significant<br>Mean difference (XBS) = 17.68; (95% CI: -1.567 to 36.92); not significant |
| o | Gaussian<br>N= 9 (control)<br>N= 9 (ABS)<br>N= 7 (XBS) | One-way ANOVA<br>Tukey test for multiple comparison<br>Control-ABS-XBS (at 2<sup>nd</sup> FE sweep) | $F = 6.779$<br>control - ABS: 95% CI: -17.09 to 39.82; not significant<br>control - XBS: 95% CI: 13.18 to 74.03; significant<br>ABS - XBS: 95% CI: 1.821 to 62.67; Not significant |
| p | Gaussian<br>N= 9 (control)<br>N= 9 (ABS)<br>N= 7 (XBS) | One-way ANOVA<br>Tukey test for multiple comparison<br>Control-ABS-XBS (at FC max) | $F = 2.381$<br>Control-ABS: 95% CI: -9.538 to 11.14; not significant<br>control-XBS: 95% CI: -2.199 to 19.90; not significant<br>ABS-XBS: 95% CI: -2.999 to 19.10; not significant |
| q | Gaussian<br>N= 9 (control)<br>N= 9 (ABS)<br>N= 7 (XBS) | One-way ANOVA<br>Tukey test for multiple comparison<br>Control-ABS-XBS (at FE end) | $F = 12.81$<br>Control-ABS: 95% CI: 23.23 to 69.40; significant<br>control-XBS: 95% CI: -5.539 to 43.82; not significant<br>ABS-XBS: 95% CI: -51.85 to -2.494; significant |
| r | Gaussian<br>N= 9 (control)<br>N= 9 (ABS)<br>N= 7 (XBS) | One-way ANOVA Tukey test<br>for multiple comparison<br>Control-ABS-XBS (at Recall) | F = 14.64<br>Control-ABS: 95% CI: 16.71 to 50.92; significant<br>Control-XBS: 95% CI: -15.21 to 21.36); not<br>significant<br>ABS-XBS: 95% CI: -49.02 to -12.46; significant |
| s | Gaussian<br>N= 9 (control)<br>N= 9 (ABS)<br>N= 7 (XBS) | One-way ANOVA<br>Tukey test for multiple<br>comparison<br>Control-ABS-XBS (at<br>Renewal) | F= 14.96<br>Control-ABS: 95% CI: .01 to 48.94; significant<br>control-XBS: 95% CI: -3.342 to 29.73; not<br>significant<br>ABS-XBS: 95% CI: -36.82 to -3.747; significant |

**Table 2:**
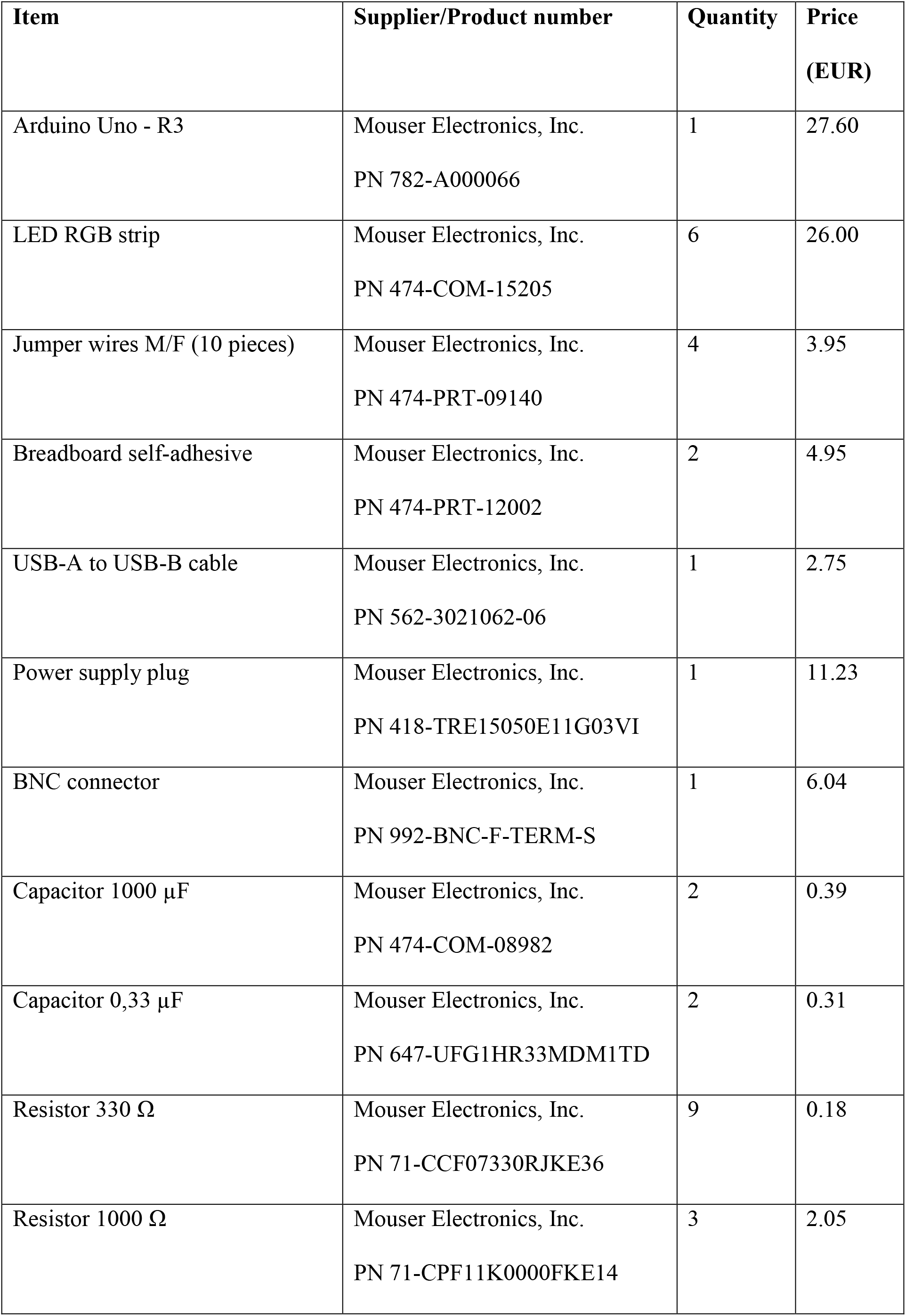
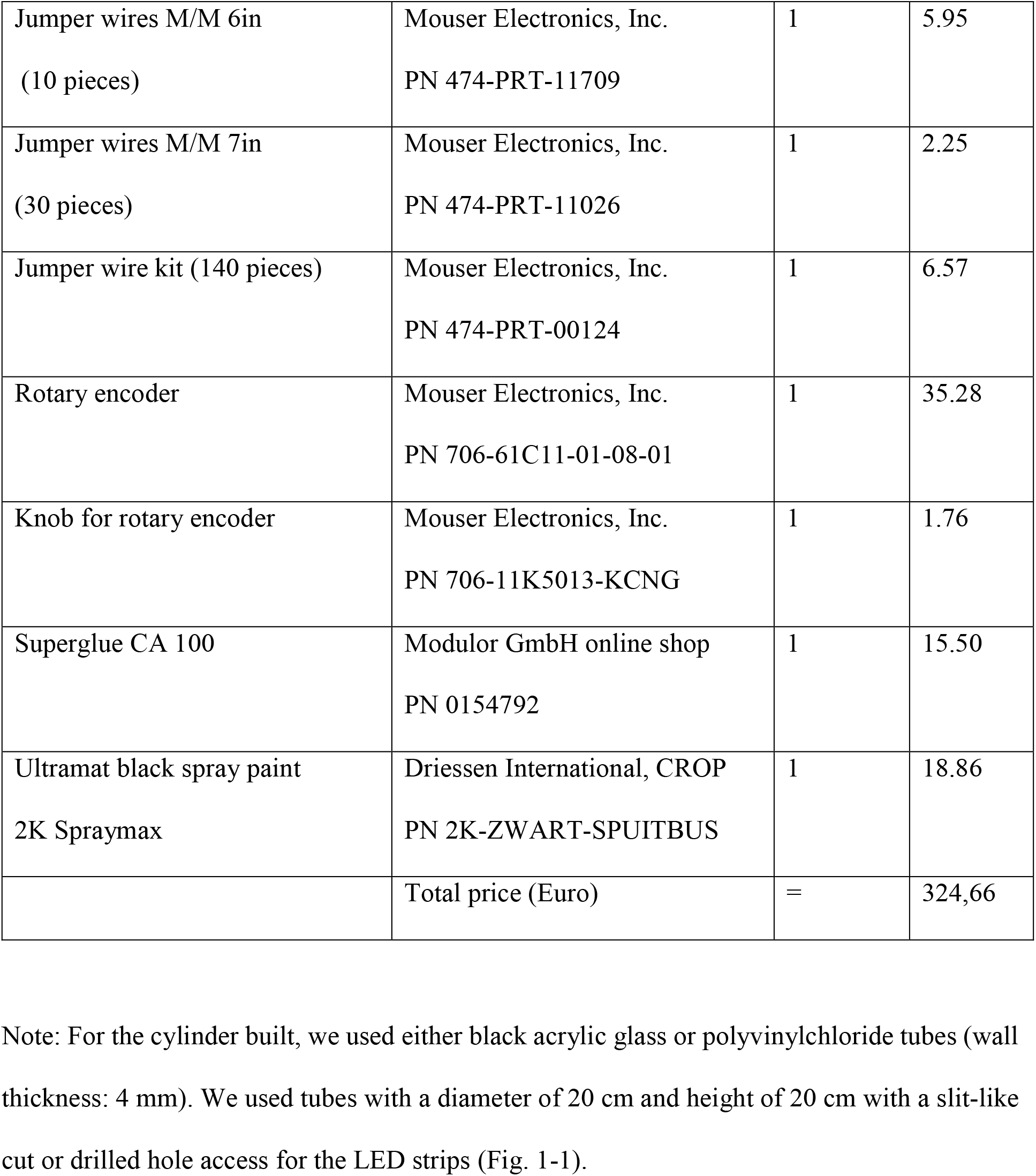
List of components necessary to build one 3MDR hardware system.

| Item | Supplier/Product number | Quantity | Price<br>(EUR) |
| --- | --- | --- | --- |
| Arduino Uno - R3 | Mouser Electronics, Inc.<br>PN 782-A000066 | 1 | 27.60 |
| LED RGB strip | Mouser Electronics, Inc.<br>PN 474-COM-15205 | 6 | 26.00 |
| Jumper wires M/F (10 pieces) | Mouser Electronics, Inc.<br>PN 474-PRT-09140 | 4 | 3.95 |
| Breadboard self-adhesive | Mouser Electronics, Inc.<br>PN 474-PRT-12002 | 2 | 4.95 |
| USB-A to USB-B cable | Mouser Electronics, Inc.<br>PN 562-3021062-06 | 1 | 2.75 |
| Power supply plug | Mouser Electronics, Inc.<br>PN 418-TRE15050E11G03VI | 1 | 11.23 |
| BNC connector | Mouser Electronics, Inc.<br>PN 992-BNC-F-TERM-S | 1 | 6.04 |
| Capacitor 1000 μF | Mouser Electronics, Inc.<br>PN 474-COM-08982 | 2 | 0.39 |
| Capacitor 0,33 μF | Mouser Electronics, Inc.<br>PN 647-UFG1HR33MDM1TD | 2 | 0.31 |
| Resistor 330 $\Omega$ | Mouser Electronics, Inc.<br>PN 71-CCF07330RJKE36 | 9 | 0.18 |
| Resistor 1000 $\Omega$ | Mouser Electronics, Inc.<br>PN 71-CPF11K0000FKE14 | 3 | 2.05 |
| Jumper wires M/M 6in<br>(10 pieces) | Mouser Electronics, Inc.<br>PN 474-PRT-11709 | 1 | 5.95 |
| Jumper wires M/M 7in<br>(30 pieces) | Mouser Electronics, Inc.<br>PN 474-PRT-11026 | 1 | 2.25 |
| Jumper wire kit (140 pieces) | Mouser Electronics, Inc.<br>PN 474-PRT-00124 | 1 | 6.57 |
| Rotary encoder | Mouser Electronics, Inc.<br>PN 706-61C11-01-08-01 | 1 | 35.28 |
| Knob for rotary encoder | Mouser Electronics, Inc.<br>PN 706-11K5013-KCNG | 1 | 1.76 |
| Superglue CA 100 | Modulor GmbH online shop<br>PN 0154792 | 1 | 15.50 |
| Ultramat black spray paint<br>2K Spraymax | Driessen International, CROP<br>PN 2K-ZWART-SPUITBUS | 1 | 18.86 |
|  | Total price (Euro) | = | 324,66 |

Note: For the cylinder built, we used either black acrylic glass or polyvinylchloride tubes (wall thickness: 4 mm). We used tubes with a diameter of 20 cm and height of 20 cm with a slit-like cut or drilled hole access for the LED strips (Fig. 1-1).

**Table 3:** List of components necessary to build the infrared floor light.

| Item | Supplier/Product number | Quantity | Price<br>(EUR) |
| --- | --- | --- | --- |
| Infrared LED | Conrad Electronic SE, Germany<br>EAN: 2050000038259 | 500 | 0.12 each |
| Europlatine breadboard | Conrad Electronic SE, Germany<br>EAN: 4016139261881 | 5 | 5.39 |
| DC supply 24V 12W | Conrad Electronic SE, Germany<br>EAN: 4024559306952 | 1 | 8.39 |
| Barrel jack adapter | Conrad Electronic SE, Germany<br>EAN: 4043619654215 | 1 | 2.24 |
|  | Total price (Euro) | = | 97,58 |

Note: As a light floor, we used white acrylic glass (1 cm) with a light transmission of <30%.

### Software accessibility

All the files for building instructions, Arduino software, bill of materials used in this study are available as Extended Data and online (https://github.com/Author_name)

## RESULTS

### Establishment of a PTSD-like phenotype of impaired fear extinction in male C57BL6 mice

Many studies showed impaired fear extinction in several mental disorders like anxiety disorders and PTSD (Shin and Liberzon, 2010). We thus set out to establish a phenotype of impaired fear extinction in male C57BL6 mice (Fig. 3A). First, we determined whether male C57BL6 mice extinguish conditioned fear in our experimental conditions. We used classic fear conditioning parameters (classic group) and showed that male C57BL6 mice both learned US-CS association and extinguished conditioned fear (paired t-test, n = 9 mice, p = 0.0^e^, Fig. 3C) for 9 days (paired t-test, n=9 mice, p=0.0^h^, Fig. 3F). We then investigated whether we can also induce a phenotype of impaired fear extinction in male C57BL6 mice (IFE group) (Myers and Davis, 2007). Despite an increase in both US intensity and the number of US-CS pairings, IFE mice expressed similar freezing responses at the end of fear conditioning compared to control mice (unpaired t-test, n=9 each, p = 0.094^d^, Fig. 3B). Yet, during fear extinction experiments, IFE mice expressed strongly impaired fear extinction (unpaired t-test, n=9 each, p=0.0136^c^, Fig. 3D). Repeated measure two-way ANOVAs revealed a significant interaction between the conditioning paradigm and the degree of fear extinction suggesting that fear extinction was significantly impaired in IFE mice throughout the time course of our extinction experiments (n=9 each, p (interaction) = 0.0301, p (time) = 0.0013, p (paradigm) = 0.0113^b^, Fig. 3B). In the initial two sweeps during the fear extinction experiments in IFE group, a fraction of mice expressed active escape behaviors during CS presentation reducing relative freezing times (Fig. 3B). IFE mice showed impaired fear extinction up to 9 days when we presented CS in a novel context (unpaired t-test, n=9 each, p=0.0628^h^, Fig. 3F), but not in a familiar context (unpaired t-test, n=9 each, p=0.894^g^, Fig. 3E). We conclude that male C57BL6 mice can express a distinct phenotype with impaired (but not disabled) fear extinction depending on specific fear conditioning parameters.

**Figure 3:**
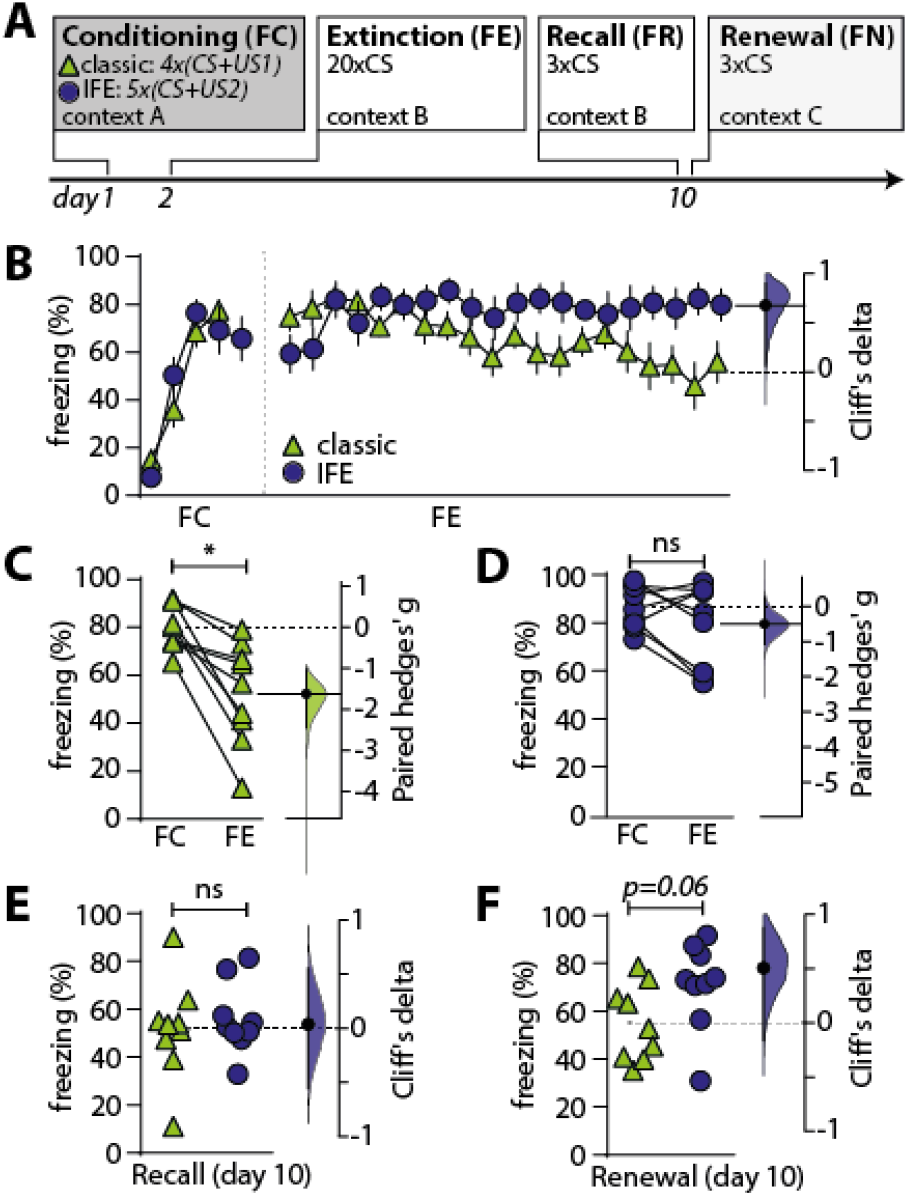
Establishment of a C57BL6 mouse phenotype with impaired fear extinction: A) Experimental timeline of fear conditioning, extinction, recall und renewal. B) Freezing behavior during FC and FE in classic (green triangle, n=9 mice) and IFE group (blue circle, n=9 mice). Mean +/-SEM for every sweep. Right floating axis depicts mean difference between classic and IFE group at FE end. C-F) Gardner-Altman estimation plots showing freezing behavior (left axis) of classic (green) and IFE mice (blue) during FC and FE (Fig. C: classic; Fig. D: IFE, paired t-test, n=9 each, p=0.151^g^), FR (E) and FN test (F) and the respective mean difference on the right axis. Asterisks indicate significant differences in two-sided permutation t-tests.

### ABS treatments persistently improve fear extinction in IFE mice

In order to validate the functionality of our 3MDR system and confirm the findings of Baek et al, we investigated whether IFE mice (control group) can persistently extinguish conditioned fear if CS presentations are repetitively paired with classic ABS stimulations (ABS group) (Fig. 4A,B). Classic ABS consisted of bilateral visual stimuli (white light) that horizontally oscillated at a frequency of 1 Hz (Fig. 4C). After fear conditioning, ABS mice showed similar freezing responses to untreated control mice (unpaired t-test, n=9 each, p=0.588^j^, Fig. 4E). During fear extinction experiments, we paired classic ABS with CS presentations and showed that coincident ABS stimulations induced a rapid and profound reduction in CS-evoked freezing behavior in comparison to untreated control mice (unpaired t-test, n=9 each, p=0.0012^k^, Fig. 4D,F). Repeated measures two-way ANOVAs revealed a significant interaction of ABS treatments with the extinction of fear suggesting that ABS mice express a strongly improved fear extinction compared with control mice (n=9 each, p(interaction)=0.0, p(time)=0.0, p(treatment) = 0.0002^i^, Fig. 4D). We conclude that coincident ABS stimulation during fear extinction is an effective way to improve fear extinction in IFE mice.

**Figure 4:**
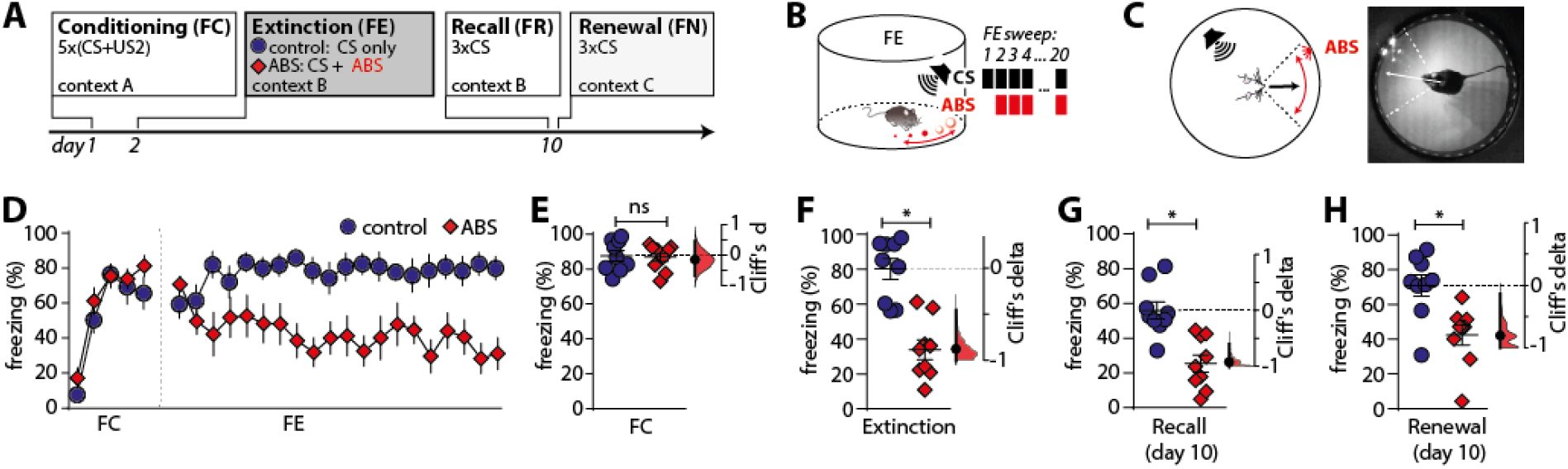
Classic ABS persistently improve fear extinction in IFE mice. A) Experimental timeline. Mice of ABS group were stimulated with classic ABS (frequency: 1 Hz, light color: white) during CS presentation in fear extinction using our 3MDR system. Mice of control group received 20 CS presentations only. B) Schematic of fear extinction in ABS group. ABS started together with the second CS presentation. C) Schematic and representative photograph showing ABS presentation in head-direction of mice during fear extinction in 3MDR cylinder. D) Freezing behavior during FC and FE in control (blue circle, n=9 mice) and ABS group (red square, n=9 mice). Mean +/-SEM for every sweep. E-H) Freezing behavior in response to CS presentation in control vs. ABS group during FC (E), FE (F), FR (G) and FN (H). Gardner-Altman estimation plots show freezing behavior (Scatter dot plot, mean+/-SEM on left axis) and the Cliff’s delta (right) between control and ABS group as mean difference and a bootstrap sampling distribution plotted on right axis. Asterisks indicate significant differences in two-sided permutation t-tests.

We then wanted to determine whether ABS reduce freezing behavior upon distraction during CS presentation or persistently improve fear extinction of IFE mice. Interestingly, ABS-treated mice expressed significantly less freezing upon CS presentation (in the absence of ABS) on day 10 in the known treatment context (unpaired t-test, n=9 each, p=0.0^l^, Fig. 4G). We conclude that ABS stimulation directly and persistently improves cue-associated fear extinction of IFE mice in the treatment context.

We then investigated whether the persistent reduction of conditioned freezing after ABS treatments depends on the contextual ABS treatment cues. Hence, we presented CS in a novel context to ABS-treated and control mice on day 10 and showed that ABS treatment improved extinction of conditioned fear even in an unknown context (unpaired t-test, n=9 each, p=0.002^m^, Fig. 4H). We conclude that ABS treatments during fear extinction significantly and persistently improved extinction of cue-associated fear in IFE mice.

### Increasing ABS brightness diminishes persistent anxiolytic effects of bilateral stimuli

We wanted to investigate whether physical stimulus properties such as stimulus brightness influence ABS-mediated anxiolytic effects. To generate extra-bright ABS (XBS), we thus increased stimulus brightness and kept all other ABS properties constant (Fig. 5A). Mice of the XBS group learned CS-US associations similar to mice of ABS and control group (One-way ANOVA with Tukey test, n=9 (control, ABS), n=7 (XBS), p=0.1158^p^, Fig. 5 B,C). During fear extinction of XBS group, we showed that XBS stimulations induced an instantaneous significant drop in freezing behavior at the first XBS-CS pairing (One-way ANOVA with Tukey test, n=9 (control, ABS), n=7 (XBS), p=0.0051°, Fig. 5B,D). Despite the strong drop of freezing at the onset of XBS-CS pairings, XBS-CS pairings induced continuously increasing freezing responses throughout fear extinction experiments compared with ABS mice (Fig. 5B). Two-way ANOVAs revealed a significant interaction between the type of visual stimulation (ABS vs. XBS) and the extinction of fear suggesting that only ABS significantly reduce conditioned fear over time (n=9 (control, ABS), n=7 (XBS), p(interaction)=0.0073, p(time)=0.0, p(treatment)=0.1095^n^, Fig. 5B).

**Figure 5:**
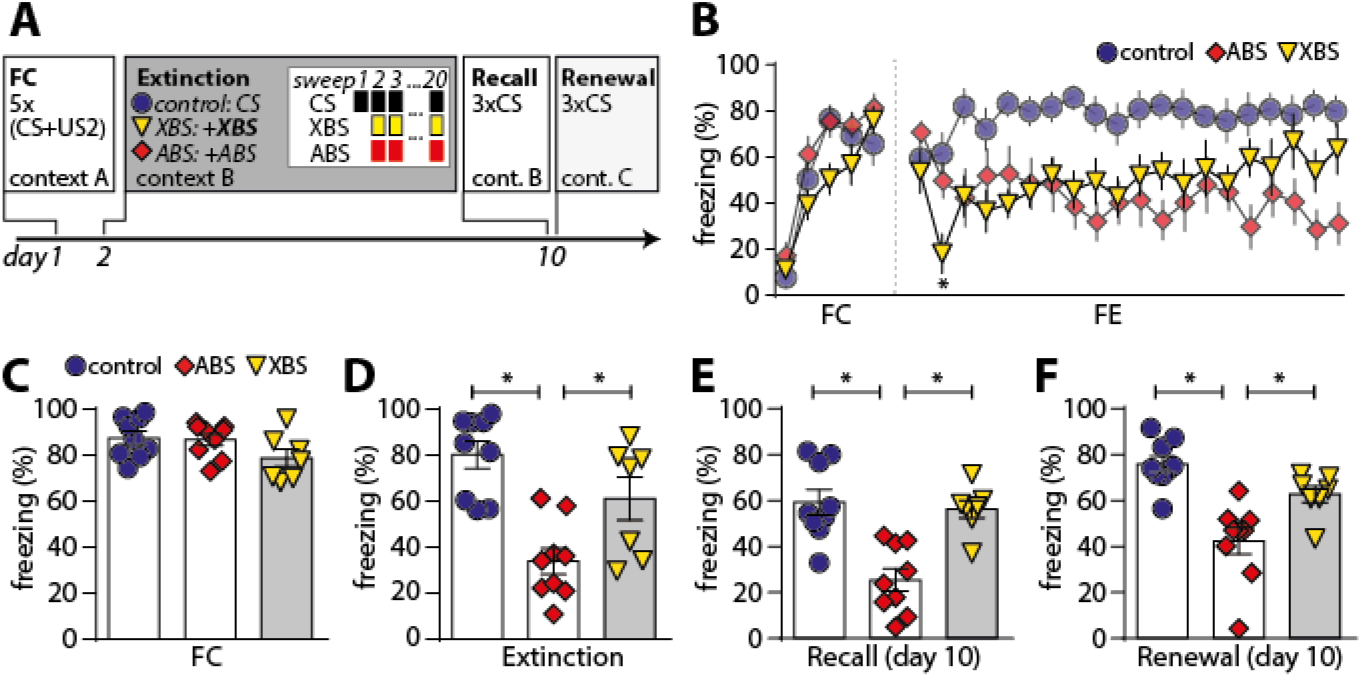
Extrabright ABS do not elicit persistent anxiolytic effects in IFE mice. A) Schematic FC and FE experiments across 10 days. During FE, we presented 20 CS like in Fig. 4/5. Alternating extrabright stimuli (XBS, yellow) were presented in the XBS group (yellow triangle) coincident to CS from the second CS presentation on. White insert in grey box shows a schematic overview of stimulus pairing in the three paradigms. For comparison, ABS (red square) and control group (blue circle) are shown. B) Freezing behavior of individual trials during fear conditioning and extinction in XBS (n=7), ABS (n=9) and control group (n=9). Mean +/-SEM. C-F) Freezing behavior in XBS, ABS and IFE groups during FC (C), extinction (D; One-way ANOVA + Tukey test, n=9 (control, ABS), n=7 (XBS), p=0.0002^q^), FR (E) and FN (F) displayed as scatter dot bar plot with mean +/-SEM. Asterisks indicate statistically significant group differences in One-way ANOVAs with Tukey’s test. Mean differences in Tukey tests: *Control (n=9) vs. ABS (n=9):* 2^nd^ FE sweep (MD=11.36; B), FC (MD=0.8000; C), FE (MD=46.32; D), FR (MD= 33.81; E), FN (MD=33.4818; F); *Control (n=9) vs. XBS (n=7):* 2^nd^ FE sweep (MD=43.61; B), FC (MD= 8.853; C), FE (MD=19.14; D), FR (MD= 3.072; E), FN (MD=13.19; F); *ABS (n=9) vs. XBS (n=7):* 2^nd^ FE sweep (MD=32.24; B), FC (MD= 8.053; C), FE (MD=-27.17; D), FR (MD=-30.74; E), FN (MD=-20.28; F)

We then wanted to determine whether XBS treatments may persistently alter extinction of conditioned fear in IFE mice in a known and unknown context - similar to ABS treatments. At day 10, we recorded freezing behavior upon CS presentation in the treatment context (recall, Fig. 5E) and in a novel context (renewal, Fig. 5F). XBS treatment (in contrast to ABS) did not persistently alter fear extinction in IFE mice on day 10 in both contexts (One-way ANOVA with Tukey test, n=9 (control, ABS), n=7 (XBS), p (recall)= 0.0^r^; p(renewal)=0.0^s^, Fig. 5E, F). We conclude that XBS treatment did not persistently improve cue-associated fear extinction in IFE mice and that ABS and XBS treatment thus do not induce the same anxiolytic effect.

## Discussion

Here, we present 3MDR, a low-cost, open-source visual stimulation system for EMDR-like interventions in mice. 3MDR standardizes the application of multiple ABS variants and the behavioral assessment in rodent EMDR-like treatments. We provide detailed hardware building instructions and open-source control software to ease the setup for inexperienced researchers. There is - to our knowledge-no commercial alternative to perform EMDR-like experiments in rodents.

Using 3MDR, we showed that classic ABS persistently improved fear extinction in mice. These results not only proved the functionality of our device but also confirmed the groundbreaking findings of Baek et al. as we showed comparable ABS-mediated anxiolytic effects. The present study primarily aimed to validate the custom-built behavioral setup, so only male mice were used as subjects. Although sex differences are of considerable interest, they are outside the scope of the present article. A second set of experiments showed that anxiolytic effects depend on physical ABS properties as bright ABS failed to persistently improve fear extinction. These different behavioral effects may be evoked by activation of different visual circuits (Williams et al., 2021), different head/ eye movements (Calancie et al., 2018; Holmes, 2019) or visual ABS features being in-/outside the mouse’ “alert range” (Yang et al., 2020). In conclusion, we show for the first time that the exact ABS composition is relevant for the behavioral effect of EDMR in mice, thereby providing a translational bridge to study the mechanisms of stimulus-specificity (Matthijssen et al., 2019) and of anxiolytic effects of ABS.

General principles of 3MDR are based on the first EMDR-like mouse model (Baek et al., 2019). The researchers showed a setup that was designed to present classic ABS to freely-moving mice during fear extinction. Baek’s setup consisted of a black treatment cylinder of 20 cm diameter inside a fear conditioning chamber in which lights are switched off.

Despite many similarities between both animal models, 3MDR offers some key advantages to Baek’s model: First, Baek et al. only used a single row of white LEDs at a fixed distance above the ground to present alternating or flickering visual stimuli. A key advantage of 3MDR is the use of programmable, multi-row RGB LED strip arrays that enables the testing of more visual stimulus characteristics such as stimulus width, color, brightness, pattern, duration, speed and direction of stimulus movement and oscillation. In particular, multi-row arrays can present ABS in vertical or angled directions, at different heights and in counter-rotation. Although an assessment of treatments with different ABS variants is of considerable interest, it is outside the scope of the present article.

Baek et al showed that ABS activate SC neurons. Yet, different SC circuits are activated depending on visual stimulus characteristics, e.g. the position within the visual field, motion direction and velocity, stimulus color and size (Cang et al., 2018; Choi and Priebe, 2020; Doykos et al., 2020; Dräger and Hubel, 1975; Ito et al., 2017; Lee et al., 2020; Li et al., 2020; Wang et al., 2015; Williams et al., 2021; Yang et al., 2020). As SC outputs and inputs are topographically mapped (Benavidez et al., 2021; Ito et al., 2017; Li et al., 2020), it is possible that ABS variants activate specific neuronal populations with different potency.

Second, Baek et al used an approach in which the experimenter chose between 1 of 4 quadrants in which ABS were presented. ABS presentation could not continuously be centered to the head direction of mice. In contrast, we use a remote control to smoothly rotate ABS in 360° inside the treatment cylinder. This rotary control can dynamically center ABS in head direction of freely-moving mice, thereby creating a more standardized situation similar to the human EMDR treatment situation. Recent studies show that mice have a sophisticated control of their visual field during freely-moving behavior (Holmgren et al., 2021; Meyer et al., 2020, 2018; Michaiel et al., 2020; Wallace et al., 2013). During attack, head movements initiate most eye movements pursuing prey (Michaiel et al., 2020). Yet, mice can also express directed saccades to a unilateral stimulus (Zahler et al., 2021). Eye movements center the cue in a small functional focus area within the visual field (Holmgren et al., 2021). Yet, it is unknown whether mice show pursuit saccades upon ABS presentation (Holmes, 2019). With respect to a highly differentiated oculomotor system in mice, it seems necessary to increase the precision of ABS presentation for future studies.

Thirdly, Baek et al recorded animal behavior in visible light. Animal videography thus contained ABS light signals, which induce strong videographic artifacts. These artifacts bias an automated pixel-change analysis towards an overestimation of ABS treatment effects as freezing during ABS is undetectable. Baek et al solved this issue by manually scoring freezing behavior during ABS treatments, yet this manual scoring (especially when ABS are visible in the video) may have big disadvantages in terms of blinded behavioral analysis. In contrast, our 3MDR system uses infrared videography in combination with a custom-made infrared floor illumination to record and semi-automatically analyze freezing behavior of mice throughout the experiments.

However, it should be noted that 3MDR uses LED strips for light generation. Although they offer advantages in terms of durability, easy setup and building costs, they are restricted to 36 pixels per row. Other solutions like flexible AMOLED displays may be favorable to generate detailed ABS in high resolution. Yet, to better understand principles of how ABS properties interact with the brain, we prioritized LED strips over high resolution displays.

It is important to highlight the versatility of our visual stimulation system for future experiments and their standardization: 3MDR has a key value to better understand the physiology how ABS interact with the brain, whether different neural circuits can be stimulated with certain ABS variants and whether ABS variants may differentially improve motivational, nociceptive or social behaviors. Together, 3MDR can not only facilitate EMDR-like interventions in mice but also future studies exploring whether visual stimuli can be used as a differential, noninvasive brain stimulation method in mice.

## ACKNOWLEDGMENTS

We gratefully acknowledge Prof. Dr. Rohini Kuner for infrastructural support and Prof. Dr. Wolfgang Eich for financial support.

## ABBREVIATIONS

3MDR: model for multimodal visual stimulation to desensitize rodents
ABS: alternating bilateral stimuli
CS: conditioned stimulus
EMDR: eye movement desensitization and reprocessing
FC: fear conditioning
FE: fear extinction
FN: fear renewal
FR: fear recall
GPIO: general purpose input/output
IFE: impaired fear extinction
IR: infrared
ITI: inter-trial interval
PTSD: post-traumatic stress disorder
SC: superior colliculus
US: unconditioned stimulus
Vis: visible
XBS: extra bright alternating bilateral stimuli

## FIGURES, EXTENDED DATA AND MULTIMEDIA

**Extended Data Figure 1-1:**
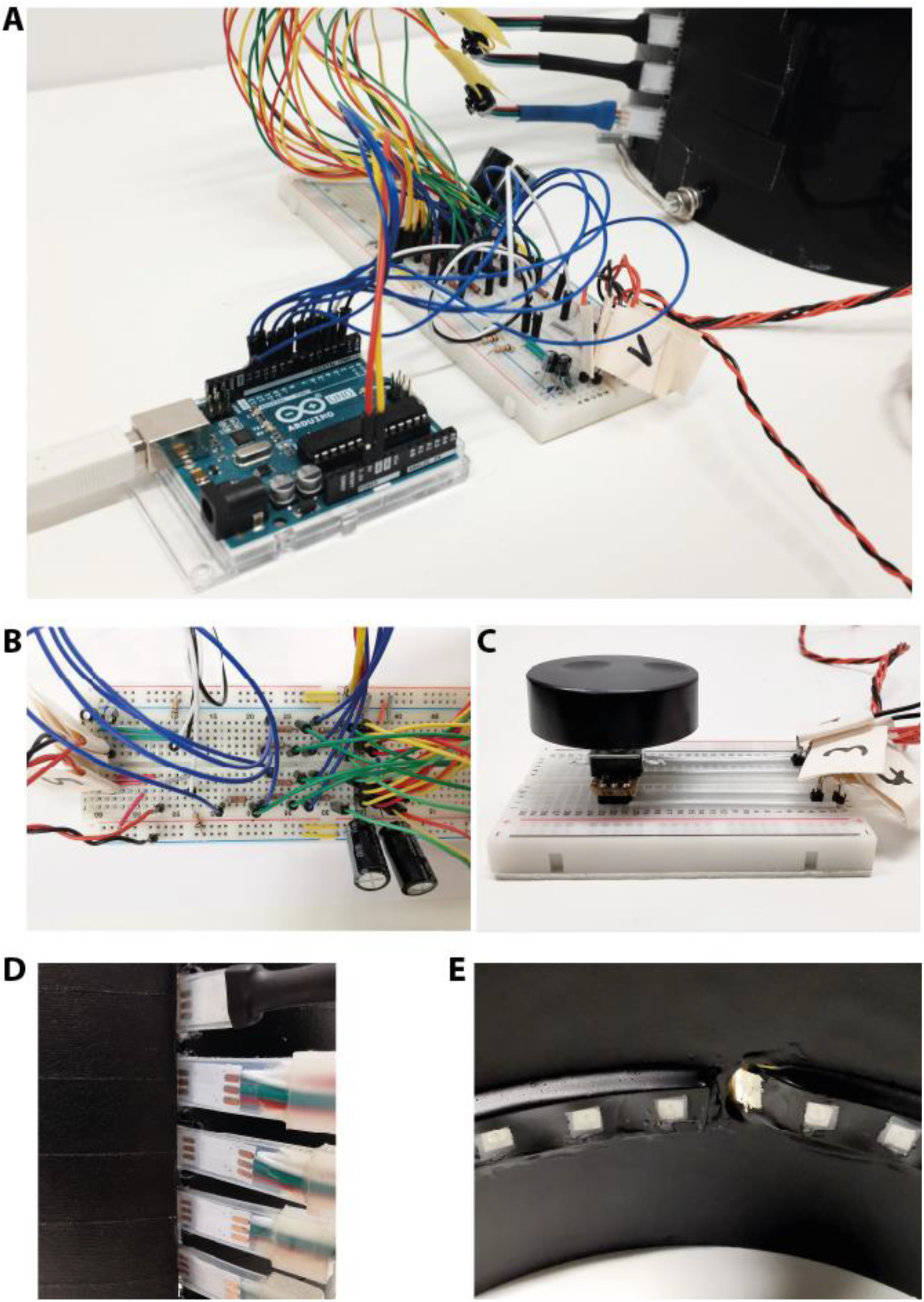
3MDR hardware. Photographs showing hardware details of 3MDR. A) Details of microcontroller und breadboard. B) Breadboard shown in high resolution. C) Remote control – rotary encoder connections. D-E) Slit- (D) and Hole-like (E) entry of LED strip to cylinder. Hole-like entry enclosed by epoxy glue.

**Extened Data Figure 2-1:**
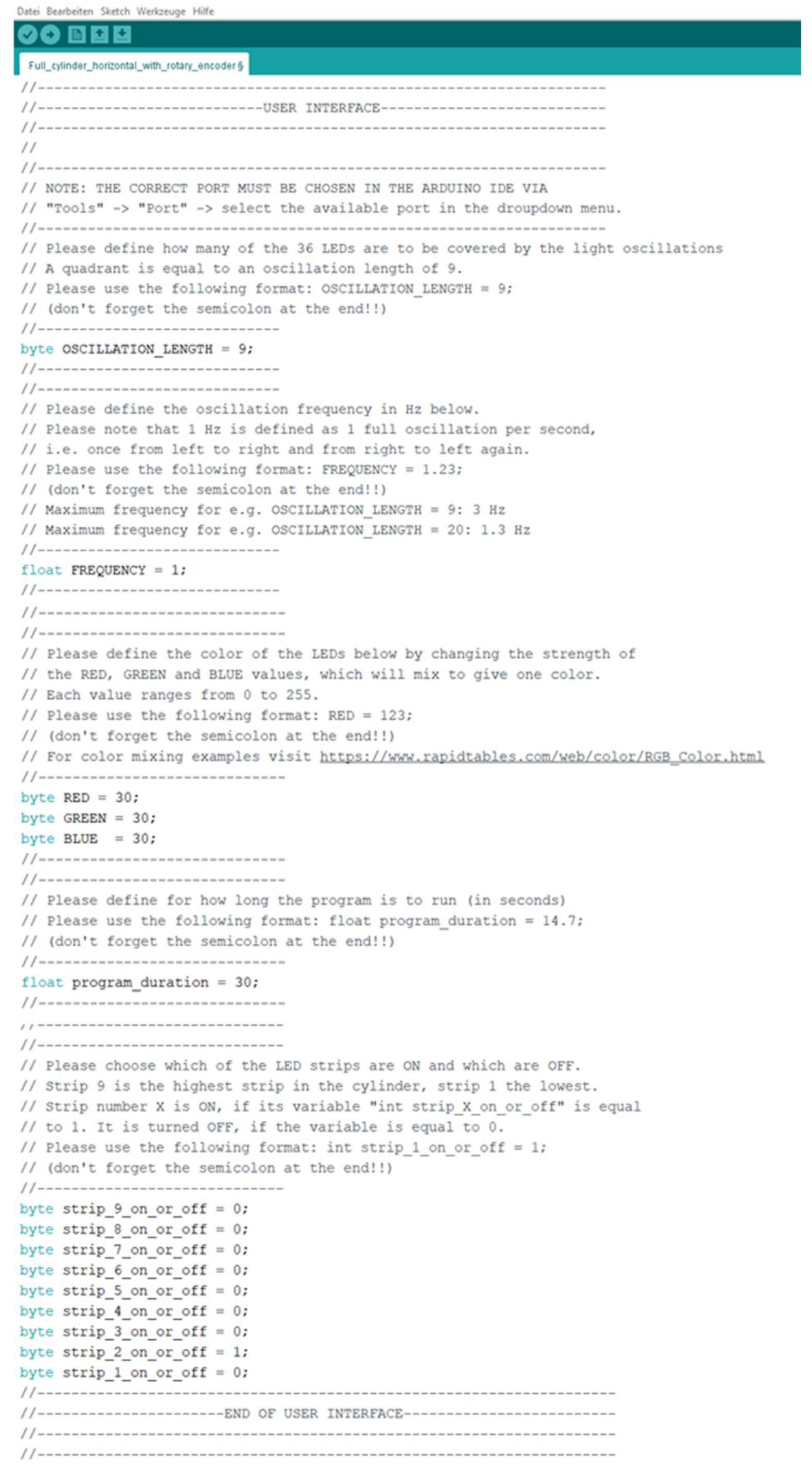
Arduino User Interface. Our Arduino User Interface that allows to adjust crucial ABS parameters. Oscillation length determines the alternation radius of the moving light. Frequency determines the speed how quick the moving light alternates in a given oscillation length. RGB color mix allows to adjust not only the color, but also ABS brightness by determining RGB values between 0 and 255. Duration determines the overall duration of the preset ABS presentation after activation. LED strip selector allows to generate different patterns, activate ABS at different absolute and relative height.

**Extended Data Figure 2-2:**
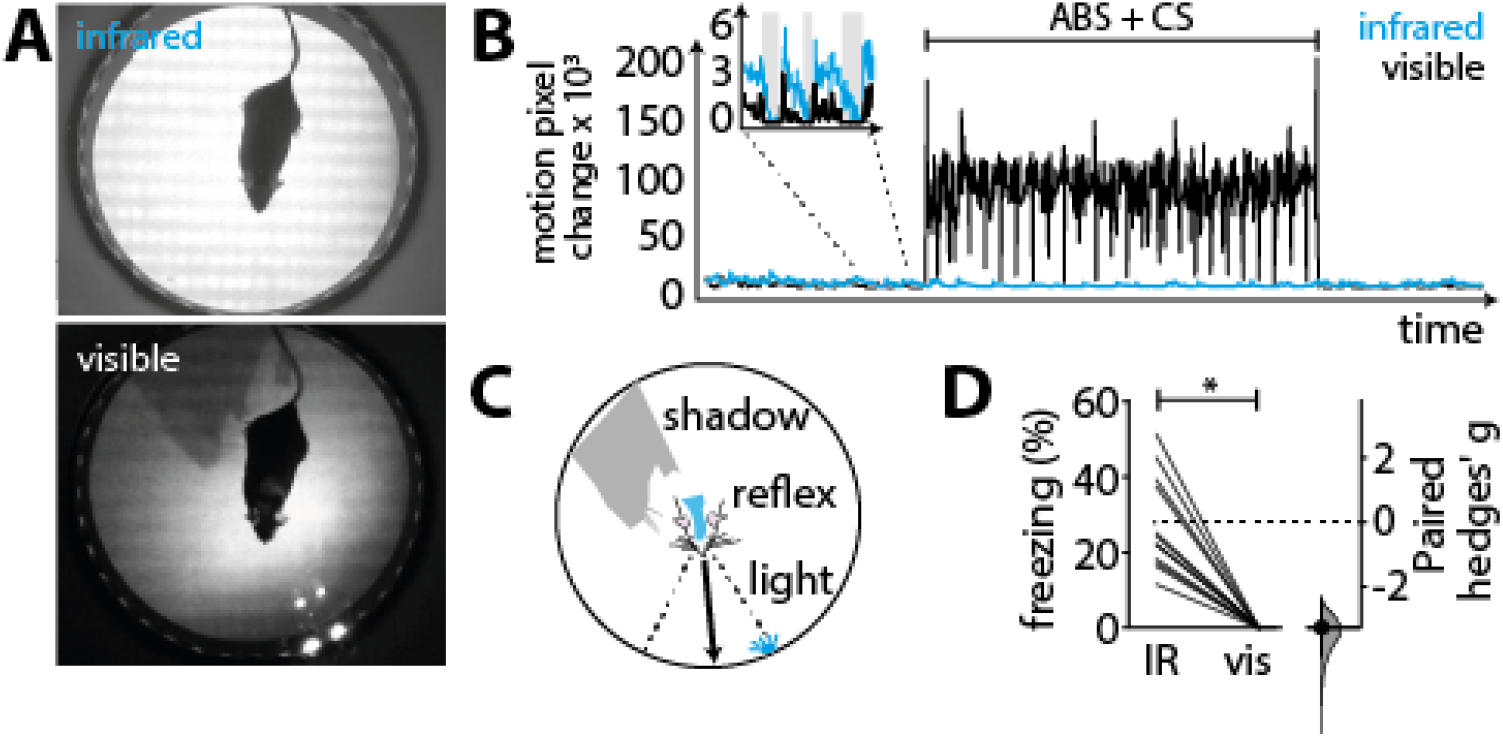
Infrared videography enables semi-automatized pixel change analysis during ABS stimulation. A) Infrared (top) versus visible light (bottom) videographic images during ABS stimulation. Using infrared videography, ABS and associated light artifacts are invisible. Infrared floor illumination prevents shadowing and enhances the contrast between animals and the background. B) Pixel change analysis on an example of parallel visible-light vs. infrared videographic recordings show massive artifacts during ABS stimulation. Insert: Without ABS stimulation, visible light and infrared videography showed comparable detection of freezing (grey bars). C) Moving shadows, fur reflexes and ABS light movements themselves distort image analysis. D) Semi-automatic freezing analysis of parallel recordings in visible versus infrared light during ABS stimulation (n=12 (2 CS per mouse, 6 mice in total). Freezing behavior shown in a slopegraph with paired Hedge’s g.

**Video 1:**
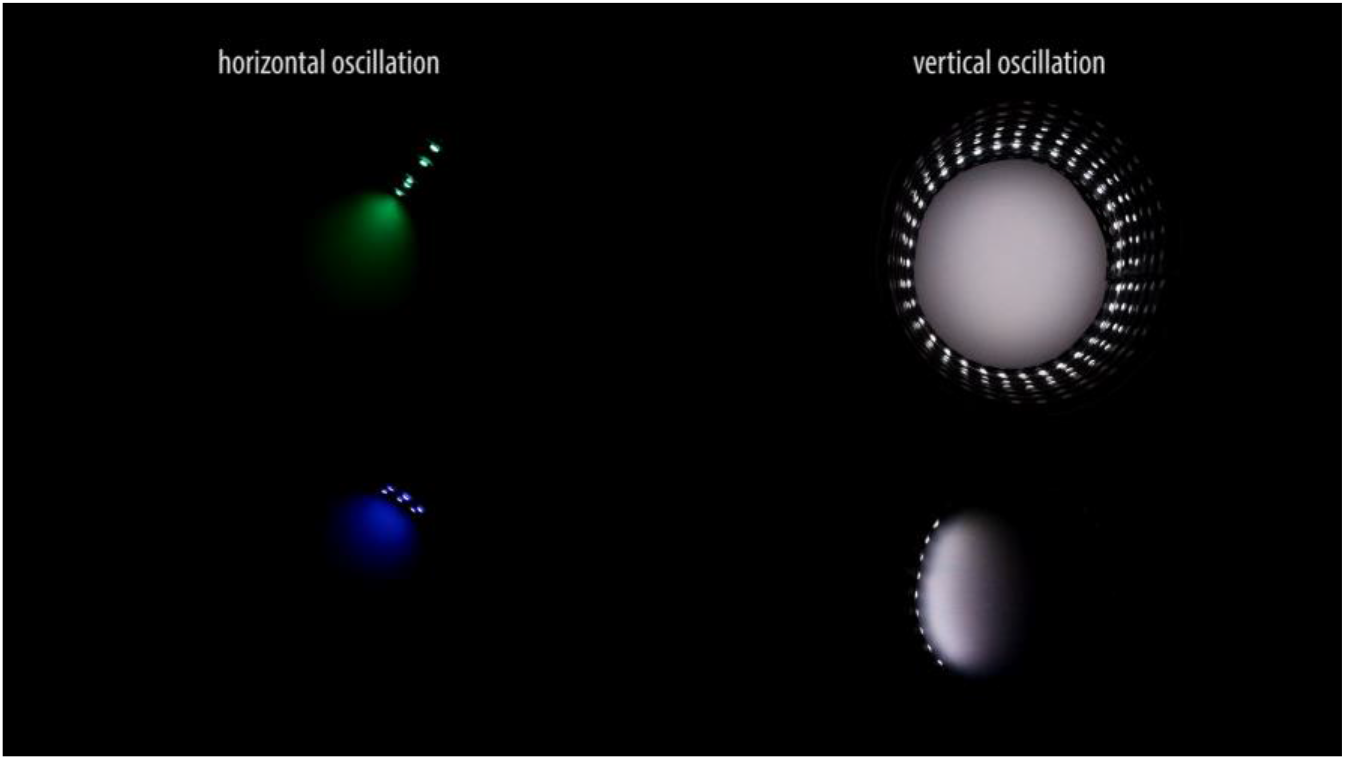
Different patterns of ABS. The video shows 4 different 3MDR oscillation patterns that can be selected and adjusted to fit the experimental requirements. Horizontally, vertically and double oscillatory or counter rotation patterns are shown. Note the variations of oscillation width, frequency, brightness and color.

**Video 2:**
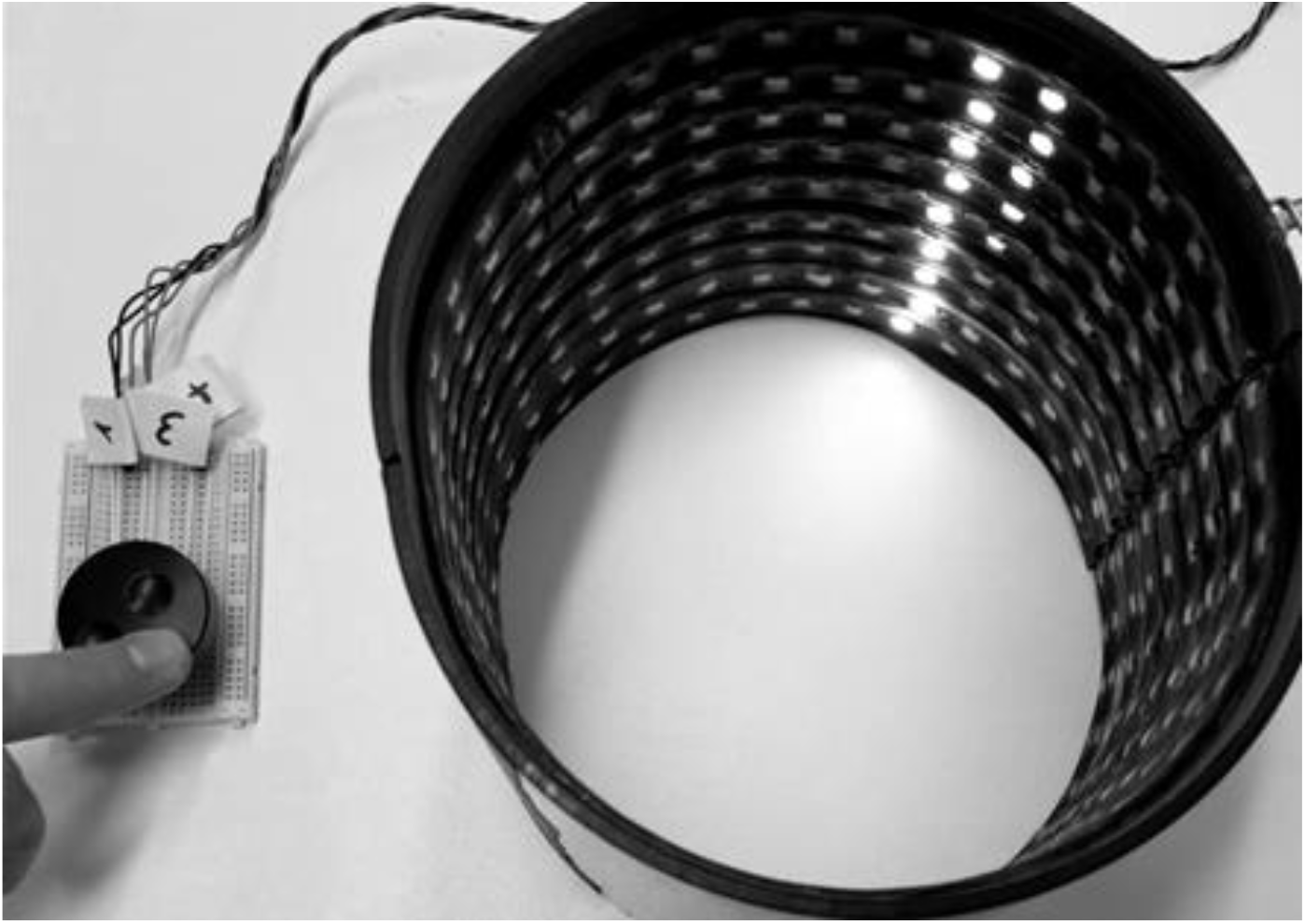
Remote control of ABS rotation inside treatment cylinder. The video shows how the 3MDR remote control rotates preset ABS inside the cylinder (first part) to center ABS in head direction of a mouse that is turning quickly in 360 degrees (second part).

**Video 3:**
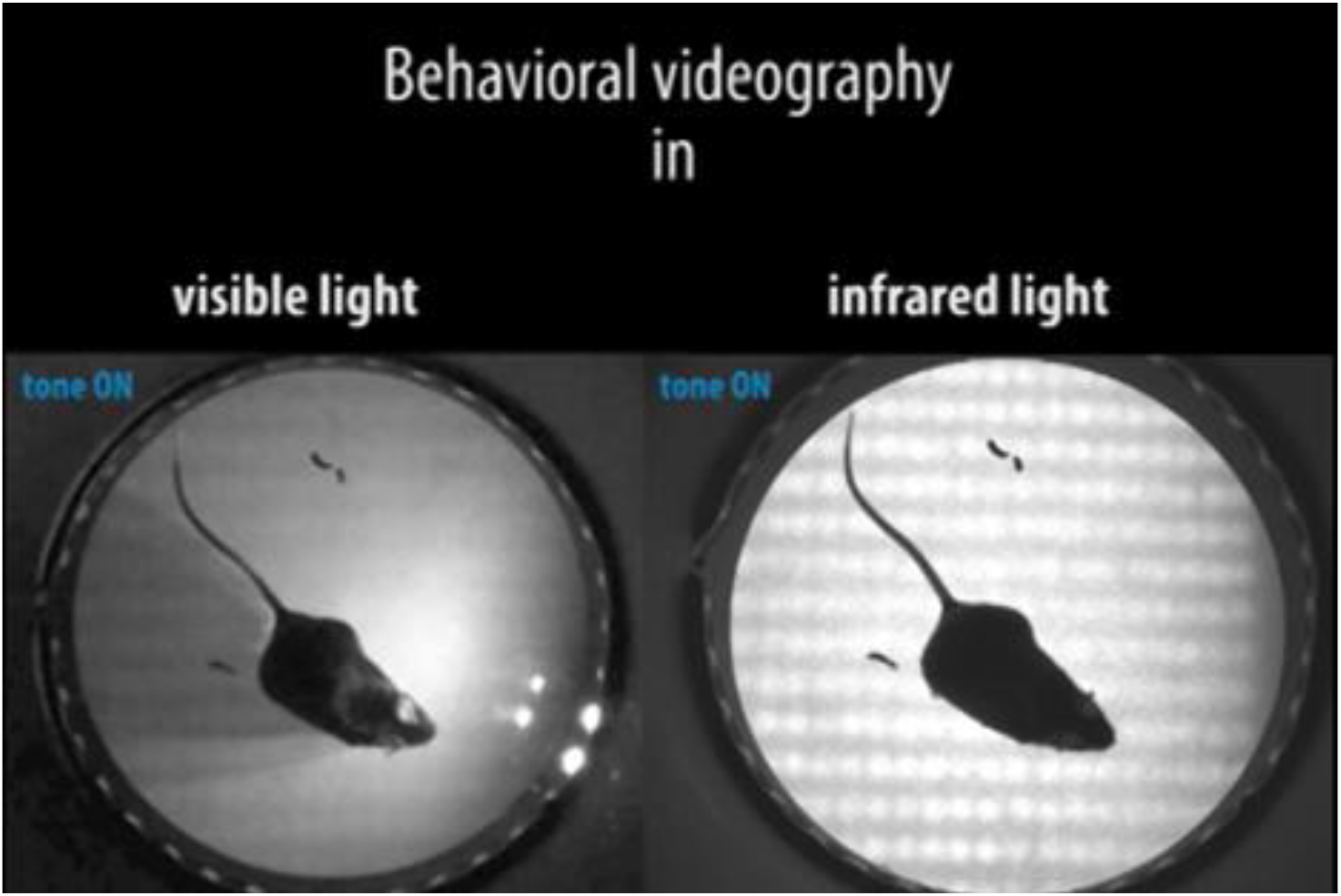
Videographic recording during ABS treatment in visible light and IR light. The video shows a parallel recording of mouse behavior before and during CS presentation and ABS stimulation in visible light and infrared light. Furthermore, the video shows that our infrared floor light increases the contrast between animals and the background.

## VISUAL ABSTRACT

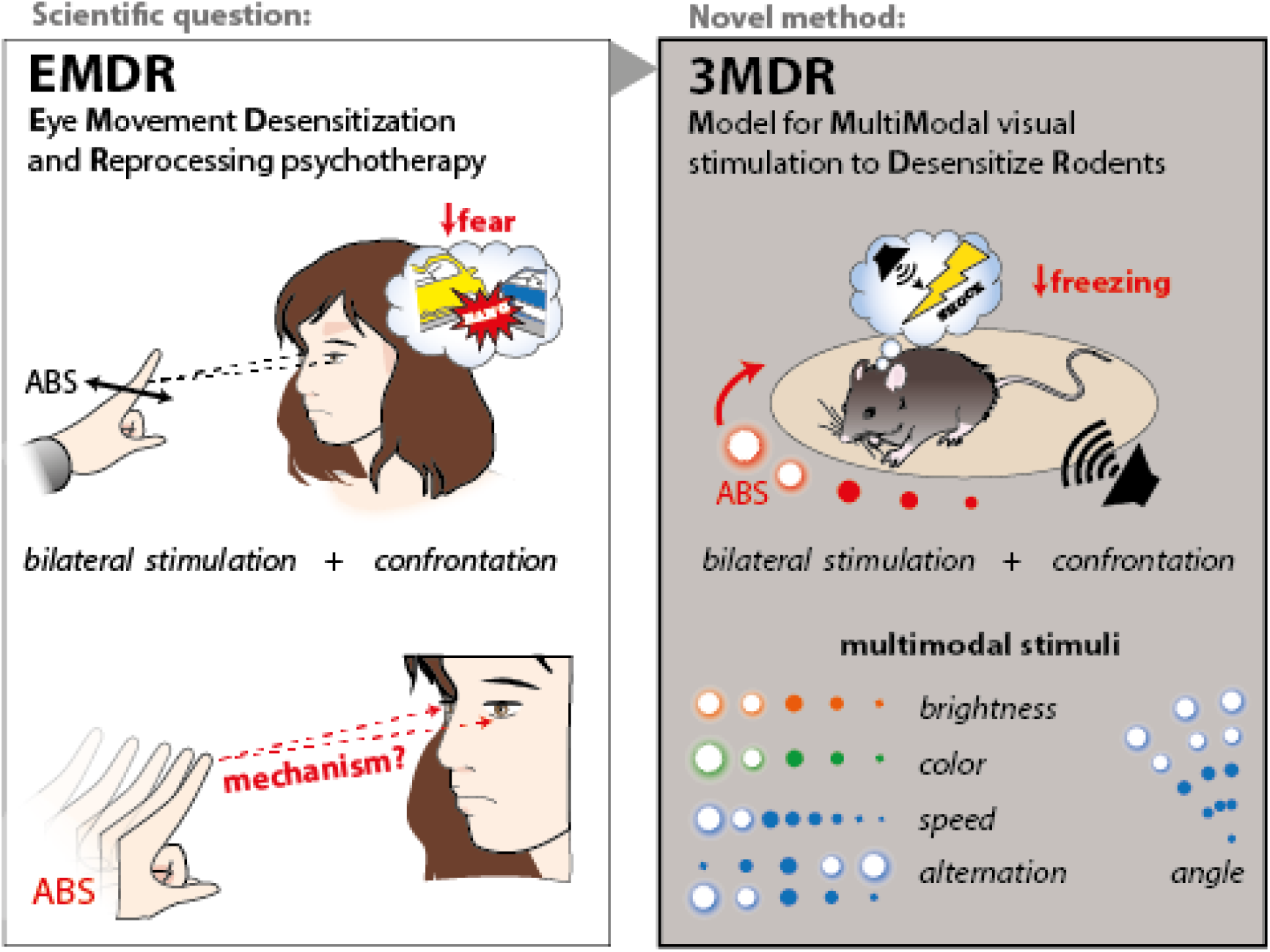

## Notes

**CONFLICT OF INTEREST:** Authors report no conflict of interest.

### Competing Interest Statement

The authors have declared no competing interest.

